# Clustering of the structures by using “snakes-&-dragons” approach, or correlation matrix as a signal

**DOI:** 10.1101/630665

**Authors:** Victor P. Andreev, Gang Liu, Jarcy Zee, Lisa Henn, Gilberto E. Flores, Robert M. Merion

## Abstract

Biological, ecological, social, and technological systems are complex structures with multiple interacting parts, often represented by networks. Correlation matrices describing interdependency of the variables in such structures provide key information for comparison and classification of such systems. Classification based on correlation matrices could supplement or improve classification based on variable values, since the former reveals similarities in system structures, while the latter relies on the similarities in system states. Importantly, this approach of clustering correlation matrices is different from clustering elements of the correlation matrices, because our goal is to compare and cluster multiple networks – not the nodes within the networks. A novel approach for clustering correlation matrices, named “snakes-&-dragons,” is introduced and illustrated by examples from neuroscience, human microbiome, and macroeconomics.

## Introduction

Inherent in our human nature is the desire to group similar objects together to better understand the world around us. It is easy to compare and group objects characterized by a single (scalar) attribute. It becomes more complex when an object is characterized by a vector of multiple attributes, although numerous clustering methods already allow for useful classifications of vectors [1]. A classification task becomes challenging with increasing complexity of the object, for example, where the interaction of object parts and attributes constitutes important characteristics of an object or a system. Indeed, some of the most engaging and challenging unresolved questions in biological and social sciences center on the comparison of functions and structures of complex systems. In this case, a system can be characterized by a matrix of interdependencies between its parts and attributes. By collecting data on the attribute levels over time or another dimension resulting in repeated measures, one can generate correlation matrices that characterize attribute interdependence and reveal important structural features of the system. In this paper, we aim to extend clustering methods to a task of comparing and classifying objects characterized by correlation matrices.

Existing methods for comparison of correlation matrices were developed mainly in evolutionary biology and applied to genetic and phenotypic variance-covariance matrices. These methods represent the differences between two matrices as one number—a similarity measure or a pairwise distance calculated by random skewers (RS), T-, or S-statistics [2–5]. Briefly, the existing methods to compare matrices are as follows: Cheverud [3] applied Pielou’s “random skewers” (RS) technique [4], which multiplies target matrices by the same randomly-generated vector (“skewer”) and averages results across numerous realizations of the vector to yield a matrix distance measure. Roff et al [2] proposed the T-method that measures the distance between matrices using a single summary statistic. More recently, Garcia proposed S-statistics, which estimates matrix distance by comparing the variance explained by the eigenvectors of each matrix [5]. These reductionist approaches have at least two limitations: (a) one number cannot adequately represent multidimensional differences; and (b) pairwise distance admits only hierarchical clustering, while other clustering methods use vectors representing multidimensional attributes of the object and might better suit the problem.

Several other approaches or variations of the above methods have also been proposed, e.g., by Goodnight and Schwartz, Calsbeek and Goodnight, Phillips and Arnold, and Flury [6–10]. However, these methods are either only applicable to a specific field of study or make strict assumptions that are not plausible in many settings. For these reasons, we focus on the distance measures from Roff et al’s T-method [2], Chevrud’s random skewers [3], and Garcia’s S-statistics [5] for comparison in the current study.

The innovative solution proposed in our paper is to create a novel although intuitively simple theoretical concept called a “snake” vector (Fig 1a), formed by making a serpentine path through the off-diagonal terms of the correlation matrix. The “snake” vector captures information on interactions between attribute variables and thus represents the system structure. Combining “snake” vectors with various other vectors representing the state of the system, e.g., vector of attribute means and variances, and overall properties of the system, e.g. number of hubs, connectedness, and small-worldness and the degree distribution [11] of the corresponding network, yields a concatenated segmental structure. We term this more complex object a “dragon” vector (Fig 1b) to designate that the analogous structure is more elaborate than the “snake”. Dragon vectors reflect not only the structural properties, but also the state of the system and allow classification based on multiple types of characterizations of complex systems. For instance, information on the initial (or average) state of the system can be described as a vector of the initial (or average) values of its attributes (creating the “head of the dragon”), while the snake formed from the correlation matrix of repeated measures will form the “tail of the dragon”. More information on the details of the snakes-&-dragons approach is provided in the Methods section. Importantly, the proposed approach allows the use of a legion of existing methods developed for clustering of multidimensional vectors.

**Fig 1.**
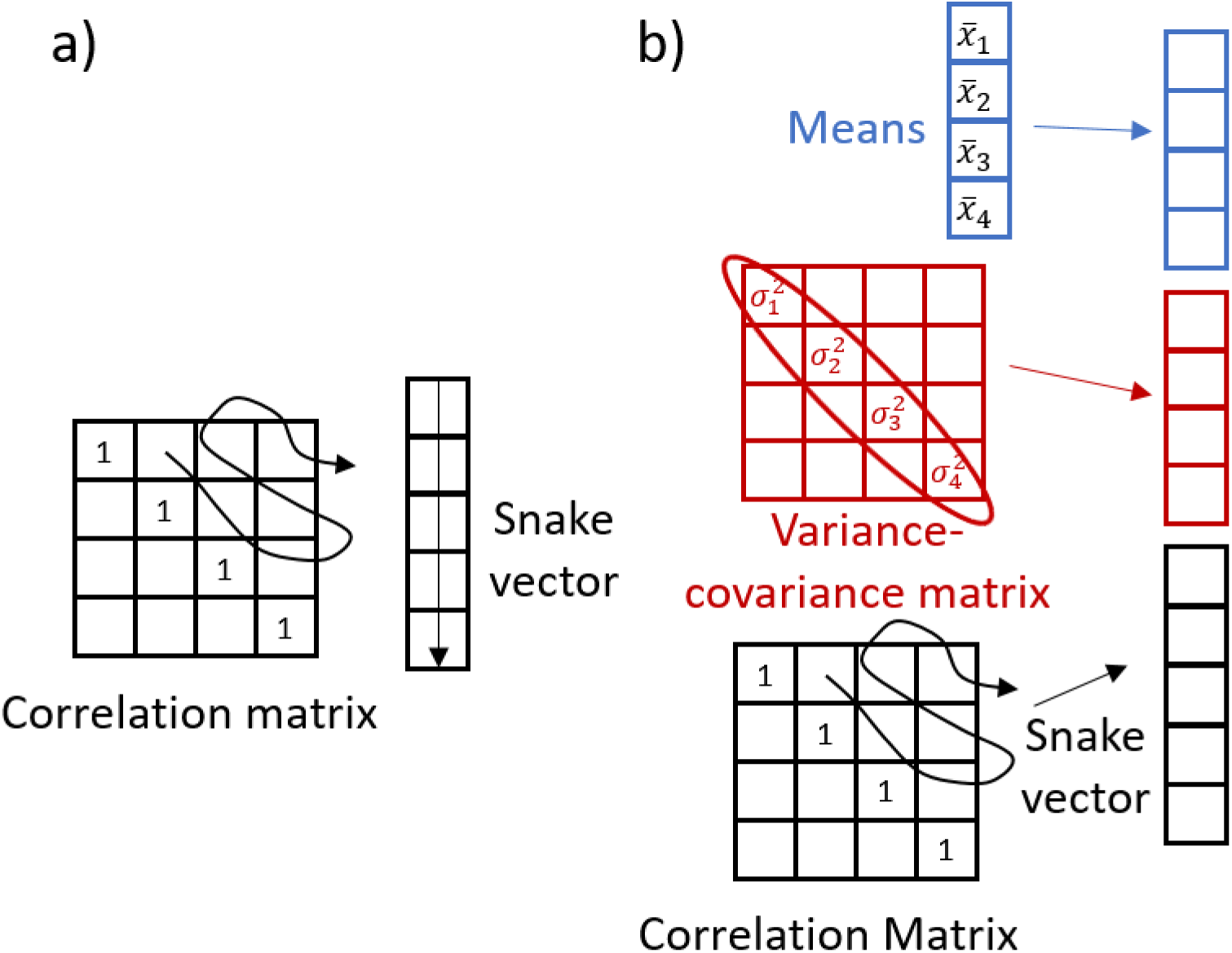
Explanation of snakes-&-dragons approach. A-snake vector. B-dragon vector. See details in the Methods section.

The proposed “snakes-&-dragons” approach is illustrated by several examples. First, we clustered brain connectivity matrices derived from resting state functional magnetic resonance imaging (fMRI) experiments [12]. Then we clustered correlation matrices describing co-occurrence of the over 10,000 microorganisms in the microbiome of gut, palm, forehead, and tongue regions of 52 students over seven weeks [13]; and finally we clustered the correlation matrices of macroeconomic development indicators from over 200 economies collected by the World Bank [14]. We clustered these correlation matrices using our proposed “snakes-&-dragons” approach and compared results with those derived from clustering based on existing measures of pairwise distances (random skewers, T- and S-statistics). We evaluated the quality of clusters by using internal validation criteria comparing within-cluster variability with between-cluster variability [15–17]. In the cases where the true cluster membership can be hypothesized, e.g., from the demographic data (for instance young vs. old), or is known as in the case of the simulated data, we determined misclassification error rates [18], and compared them using our and other approaches. Next, we examined the number of significantly different variables across the clusters, testing all the variables used for clustering and other variables such as demographics. This provides not only the proof of cluster distinctiveness but also the information about the possible factors driving cluster membership. We believe that the high values of cluster validation criteria together with the high percentage of significantly different variables across the clusters could illustrate that identified clusters meet the concise definition of clustering given by Liao [19] as: “identifying structure in an unlabeled data set by objectively organizing data into homogeneous groups where the within-group-object dissimilarity is minimized and the between-group-object dissimilarity is maximized.”

## Materials and methods

### Data sets

First, we briefly describe data sets used to illustrate and validate our proposed snakes-&- dragons approach to clustering correlation matrices.

#### Brain connectivity matrices from old and young healthy subjects

Brain connectivity matrices arise from the observation that the blood oxygen level-dependent (BOLD) fMRI signal is correlated between spatially separated but functionally related brain regions [20-21-]. Multiple fMRI studies of resting state brain activity showed that matrices of correlation coefficients of BOLD signal between brain regions (connectivity matrices) differ in health and disease, especially in mental disorders [21–22]. Several studies demonstrated changes in brain connectivity matrices related to aging [23–24]. A pilot data set of brain connectivity matrices used in our study was created at Washington University in St. Louis. It includes connectivity matrices from 20 healthy subjects older than 60 (#1- #20) and 17 subjects younger than 27 (#21- #37). The data set of older subjects was obtained with permission from the Washington University Alzheimer’s Disease Research Center and served in their study as a control group (Clinical Dementia Rating = 0 and CSF biomarker negative). The data set of younger subjects is the same as used in [25–26] with mean age 23.1 years and range 18-27; all of them were screened to exclude neurological impairment and use of psychotropic medications. Connectivity matrices with 36 functional areas were then calculated from the fMRI scans using the Washington University pipeline described in [27]. Then the 37 connectivity matrices were clustered by using our snake vector approach, without using any demographic information.

#### Brain connectivity matrices from the Brain Genomics Superstruct Project

The Brain Genomics Superstruct Project Open Access Data Release (GSP) is a carefully vetted collection of neuroimaging, behavior, cognitive, and personality data for 1,570 human participants (ages 18-35)[12]. GSP data include not only demographic data (age, handedness, sex) for all participants, but also anatomical information on the brain and its regions for each of participants. The 169 brain areas were divided into 10 networks: visual foveal (VFN), visual peripheral (VPN), dorsal attention (DAN), motor (MN), auditory (AN), cingulo-opercular (CON), ventral attention (VAN), language (LN), fronto-parietal (FPN), and default mode (DMN) [26]. Connectivity matrices were calculated from the fMRI scans using the Washington University pipeline [27] for the first 500 participants of the GSP cohort that had two BOLD fMRI runs and cognitive behavioral data.

#### Microbiome data for healthy college-age adults

Flores et al collected longitudinal (10 weeks) data to analyze temporal dynamics of forehead, gut, palm, and tongue microbial communities among 85 healthy college-age adults from three US universities [13]. A 49-question demographic, lifestyle, and hygiene survey augmented the weekly sample collection. Based on relative abundance of over 10,000 microbial species measured as operational taxonomic units (OTUs) in each sample, investigators found high variability in the microbiome over time. In our study, we aim to characterize the temporal changes in the microbiome by exploring correlations between weekly samples of microbiomes within each individual. By clustering individuals’ correlation matrices, we identified subgroups of students representing different patterns of microbiome dynamics.

#### Macroeconomics development indicators from the World Bank

Since 1960, the World Bank has collected 1,500 yearly macroeconomic development indicators from over 200 economies, including: 1) gross domestic product (GDP), 2) unemployment, 3) inflation, 4) net trade in goods, 5) labor force participation, 6) foreign direct investment, and 7) gross domestic savings [14]. As a proof-of-concept example, we used the time series data on the seven indicators to create 7-by-7 correlation matrices for each of the 200 economies and then clustered them by using snake vectors.

### Analytical methods

In this paper, we compare and cluster correlation matrices from the above four data sets by using existing methods for matrix comparison and our novel “snakes-&-dragons” approach.

#### Existing methods to compare matrices: random skewers, T-statistic, S-statistic

Approaches to compare and calculate distances between matrices were developed in evolutionary biology and might be unfamiliar to researchers outside of that field. Therefore, we briefly describe three of the existing approaches used in this paper: random skewers (RS), T-statistic, and S-statistics. The RS procedure samples from a uniform [−1, 1] distribution to form random vectors [28]. Multiplying correlation matrices by these vectors yields response vectors. If the compared correlation matrices are similar, the responses to the same selection vector should be similar as well. The correlation among response vectors is averaged over multiple random vectors—100 replicates in our example—to estimate similarity between two objects. Another method for comparing matrices is the T-statistic [2], describing dissimilarity between two matrices as the sum of the absolute differences between corresponding matrix elements. The third method is the so-called S-statistic [5]. Garcia introduced three S-statistics to represent the divergence between two correlation matrices, all based on the idea that if two covariance matrices are similar, an eigenvector set resulting from principal component analysis (PCA) of one matrix will explain a similar amount of variation in the other matrix. We considered the first, S1, which Garcia described as a general measure of differentiation, characterizing the ability of eigenvectors from one sample to explain the variation in the other sample. By contrast, S2 compares orientation of eigenvectors of the same ordinal position in the two sets and S3 evaluates differences in shape of eigenvectors in the same ordinal position between the two sets. We performed hierarchical clustering based on the resulting similarity matrices.

#### Creating “snakes-&-dragons”

We propose to extract details from correlation matrices into a new object that we call a “snake” vector. The “snake” vector forms from a serpentine path through the off-diagonal terms of a correlation matrix and captures information on interactions of the variables, i.e., the system structure (Fig 1a). Many methods exist for clustering of vectors, allowing for the choice of the optimal clustering method for a given data set or problem. To augment and complement the information on the structure of the systems with the information on the state of the systems, we additionally introduce the class of objects that we call “dragon vectors” or “dragons”. Here we suggest four types of dragons. Dragon 1 integrates state descriptors and structural descriptors by concatenating the snake vector with a vector of variable means and a vector of variable variances (Fig 2a). Dragon 2 (Fig 2b) integrates structural descriptors with overall network property information. While the snake vector contains individual correlations between system attributes or between nodes of a network to represent structural descriptors, measures of network integration can describe the system in a different way. For example, average connectivity, number of nodes/hubs, average or shortest path length, or number of first neighbors have previously been used to characterize networks [11, 29-30]. These measures can be concatenated with the snake vector to form a dragon for clustering. Dragon 3 (Fig 2c) is created by combining correlations along multiple dimensions or locations. We used this approach in the analysis of the microbiome data set, which contains measures of microbial OTUs at four sites on the human body at several time points in many subjects. The correlation matrix for each body site yields a different snake vector. By concatenating multiple snakes, all data descriptions can influence the clustering. Similarly, Dragon 4 (Fig 2d) can be created by combining different types of data, e.g., correlation matrices of clinical, transcriptomic, proteomic, and metabolomic variables derived from repeated measures combined with the genomics data and baseline demographics and clinical data, which would create the “head” of the “dragon”. While snake vectors can be clustered as they are, since the elements of the correlation matrices are always in the range from −1 to 1, dragon vectors require several refinements prior to processing. First, clustering algorithms often gravitate toward elements of greater magnitude. We thus put all variables on a common scale to ensure all variables can fairly influence the decision-making. When a data set has a natural comparison group, e.g., with cases and controls, observations on cases can be centered and scaled using the mean and standard deviation of the corresponding variable among controls. In the absence of such a control group, as in this study, we center variables by each variable’s mean and scale by the square root of its average variance. Additionally, cluster results should not be affected by including variables reflecting redundant information. To mitigate that prospect, we suggest performing PCA on the matrix of assembled dragon vectors and then clustering based on the principal components.

**Fig 2.**
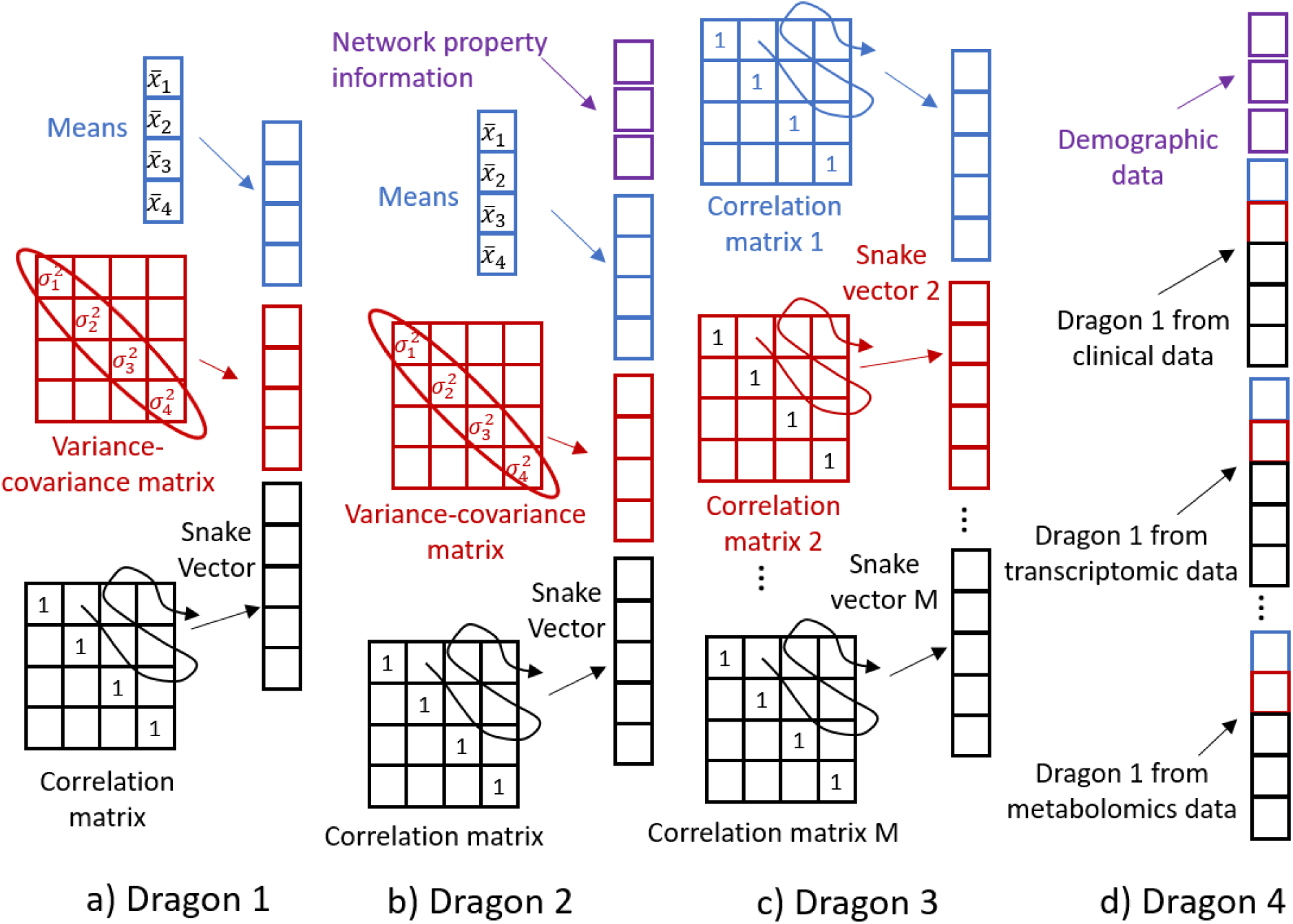
Four types of dragon vectors. A-Dragon 1, includes means and variances of the variables. B-Dragon 2, includes also overall network property information. C- Dragon 3, combines correlations along multiple dimensions of the data matrix or multiple locations. D-Dragon 4 is composed of several dragons presenting different types of clinical and omics data.

#### Clustering methods

Many clustering methods exist, including k-means clustering, fuzzy k-means clustering, hierarchical clustering, k-medoids, affinity propagation, and others [1]. Choosing among algorithms and choosing the number of clusters is often achieved using internal validation statistics, such as Calinski, silhouette, or connectivity [15–16]. None of the clustering methods is ideal in all settings, and the optimal choice depends on the underlying data’s properties, which is not always recognized by the users of clustering algorithms. For example, Dolnicar found that clustering studies typically do not match data conditions with clustering methodology, but instead just use Ward’s hierarchical and k-means clustering [31]. Halkidi et al noted that many studies omit cluster validation, despite its importance and the availability of tools for implementation [17]. They suggested that new clustering algorithm development should include simulated data sets that mimic the properties of biological data to allow for controlled study of an algorithm’s sensitivity. Our group recently compared three clustering methods—hierarchical, k-means, and k-medoids—using simulated targeted proteomics data [18]. We demonstrated that k-means had the lowest misclassification error for identifying biomarker signatures, but also that results varied with different correlations between biomarker levels. The study illuminated the importance of the structure of the correlation matrix of the variables in determining the optimal clustering method [18].

Clustering in this study is performed using a resampling-based consensus clustering method introduced by Monti et al [32]. As implemented in our study, this method can be briefly described as follows. We performed 1,000 instances of random samplings with replacement, each selecting a subset including 80% of N objects (snake or dragon vectors under study). We then partitioned each of the subsets into clusters using a k-means clustering algorithm (implemented as the MATLAB® function kmeans; MathWorks, Natick, MA) with k value scanned from 2 to 8. Then the N x N consensus matrix was created representing the results of these 1,000 partitions. Each element of the matrix represented the proportion of times that the two objects were included in the same cluster, i.e., the ratio of the number of times a given pair of objects were included in the same cluster to the number of times both of the objects were selected in the random 80% subset. Therefore, each element of the matrix can be interpreted as a probability that two objects belong to the same cluster. Hierarchical clustering (using MATLAB® function clustergram) was then performed using elements of the consensus matrix as the distance measure between objects. Resulting clusters (for each scanned value of k) were then examined by using Calinski’s “quality of clustering” criterion, which compared the between-cluster differences with the within-cluster differences and allowed determination of the optimal number of clusters [15].

For RS, T-, and S- statistics, hierarchical clustering was used since it is the only method that can work with these measures of pairwise distances between objects (vectors, matrices). Hierarchical clustering was performed using the clustergram MATLAB function with the Ward distance option.

#### Simulating correlation matrices with a controlled noise level

When working with real data, one disadvantage is that the true cluster membership is not known, so it might be difficult to evaluate the misclassification error rate. Thus, in order to evaluate clustering of connectivity matrices using snake vectors, we created simulated data that had clear “labels” (e.g., older or younger brain connectivity matrices). In this study, we selected two substantially different brain connectivity matrices, #1 and #29, as representatives of old and young brains, respectively (from the 37 healthy young and old subjects pilot data set described above). Based on these two prototype matrices, we simulated two matrix classes by adding a controlled amount of noise. Since correlation matrices need to satisfy certain conditions (i.e., being a positive-semidefinite matrix), we cannot just add noise to each component of the matrix. Instead, we used the procedure suggested by Schafer et al, which simulates noise by repeatedly sampling from multivariate normal distributions with given standardized covariance matrices [33]. Briefly: we take the *q x q* brain connectivity matrix and use it as a covariance matrix to simulate the multivariate normal distribution from which we sample n times to generate a *q x n* data matrix. Then, we calculate the *q x q* correlation matrix from this data matrix. The higher the *n* the closer the new correlation matrix to the original connectivity matrix will be. Decreasing *n* may be viewed as adding noise, since the role of randomness is higher when the normal distribution is sampled more sparsely. This procedure allows the amount of noise to vary by changing a *q/n* ratio, where *q* is the number of variables (here number of brain regions *q* =36) and *n* the number of times the multivariate normal distribution is sampled to create a data matrix used to calculate the correlation matrix. Importantly, each time we randomly sample the multivariate normal distribution, we get a different *q x n* data matrix and the *q x q* correlation matrix, even for the same value of *n*. Fig 3 shows single instances of simulated correlation matrices when the *q/n* ratio is set to 0.1, 3, 6, 9, and 12 for brain connectivity matrix #1. The similarity of the simulated matrices with the original prototypic connectivity matrix #1 is clearly decreasing.

**Fig 3.**
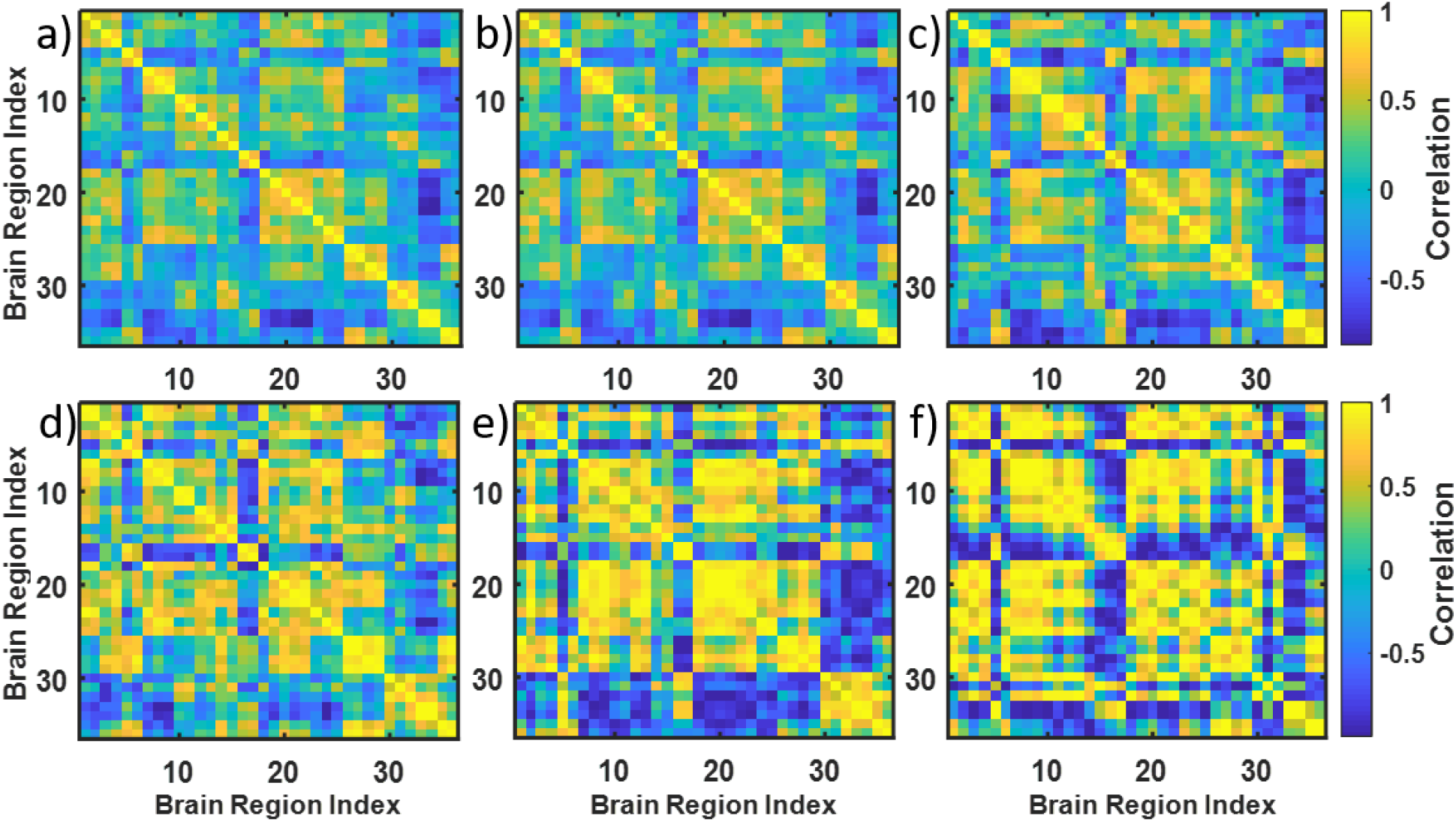
Simulating connectivity matrices with increased noise level. A-original matrix #1. B – simulated matrix with *q/n*=0.1, *n*=360; C – *q/n*=3, *n*=12; D- *q/n*=6, *n*=6; E-*q/n*=9, *n*=4; F- *q/n*=12, *n*=3.

Figs 4a-b demonstrate how the instances of simulated correlation matrices differ from each other for given values of *n*. As seen, the variability across the instances is higher the lower the *n*. To test and compare the performance of the snake vector approach with the existing measures of matrix dissimilarity, we simulated 20 such matrices for each value of the *q/n* ratio for prototypic old and prototypic young brain connectivity matrices (#1 and #29) and conducted clustering on the 40 simulated connectivity matrices for each *q/n* value. This enabled us to compare the ability of the various clustering methods to correctly classify the correlation matrices as young or old in the presence of an increased level of noise. To make better sense of what *q/n* means in terms of added noise and variability of the simulated connectivity matrices, we calculated the histograms of standard deviations of the elements of the simulated connectivity matrices for various *q/n* values (shown in Fig 5a). Clearly, standard deviations are higher for larger *q/n* values. Then, we defined the signal/noise ratio (SNR) describing difference between two clusters of correlation matrices as follows:

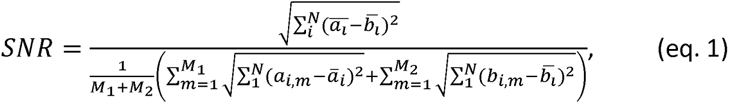

where *N* is the length of snake vectors, *ā_l_* is the *i*-th element of the average snake vector for cluster 1, *b̄_i_* is the *i*-th element of the average snake vector for cluster 2, *M*_1_ is the number of simulated matrices in cluster 1, *M*_2_ is the number of simulated matrices in cluster 2, *a_i,m_* is the *i*-th element in the snake vector obtained from *m*-th simulated correlation matrix and *b_i,m_* is the *i*-th element in the snake vector obtained from *m*-th simulated correlation matrix. Note that the numerator in eq. 1 is the Euclidian distance between the centroids of the two clusters, which is equal to the distance between the snake vectors of the prototypic connectivity matrices, while the denominator is the measure of the average within cluster Euclidian distances. Fig 5b demonstrates how SNR defined by (eq. 1) depends on the *q/n* value.

**Fig 4.**
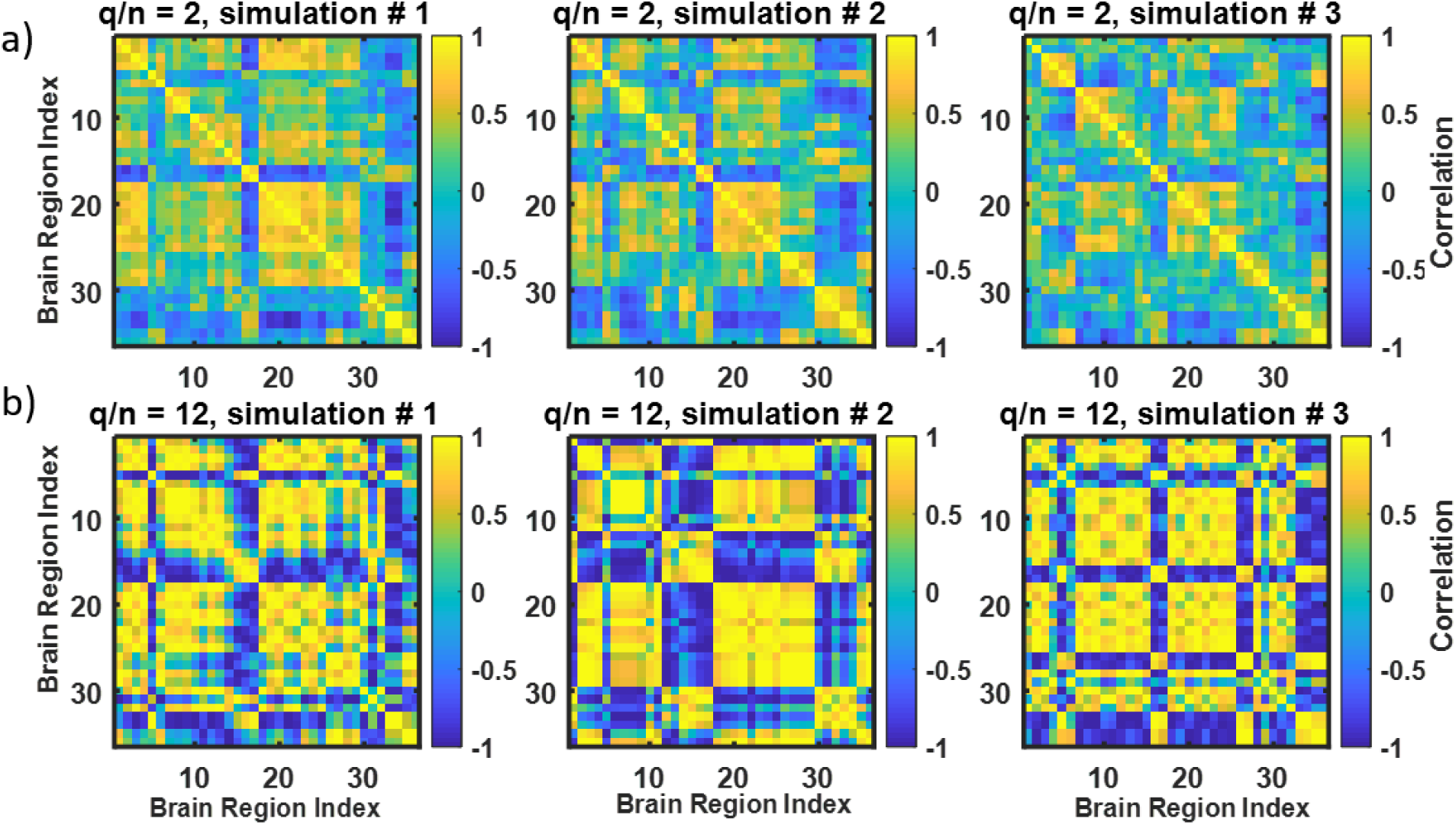
Increased variability of simulated correlation matrices with increased *q/n* value. A-3 instances of correlation matrices generated from the connectivity matrix #1 using *q/n*=2, *n*=18; *B*-3 instances of correlation matrices generated from the connectivity matrix #1 using *q/n*=12, *n*=3. See how variability of the matrices is increased in B (*q/n=12*) versus A (*q/n=2*).

**Fig 5.**
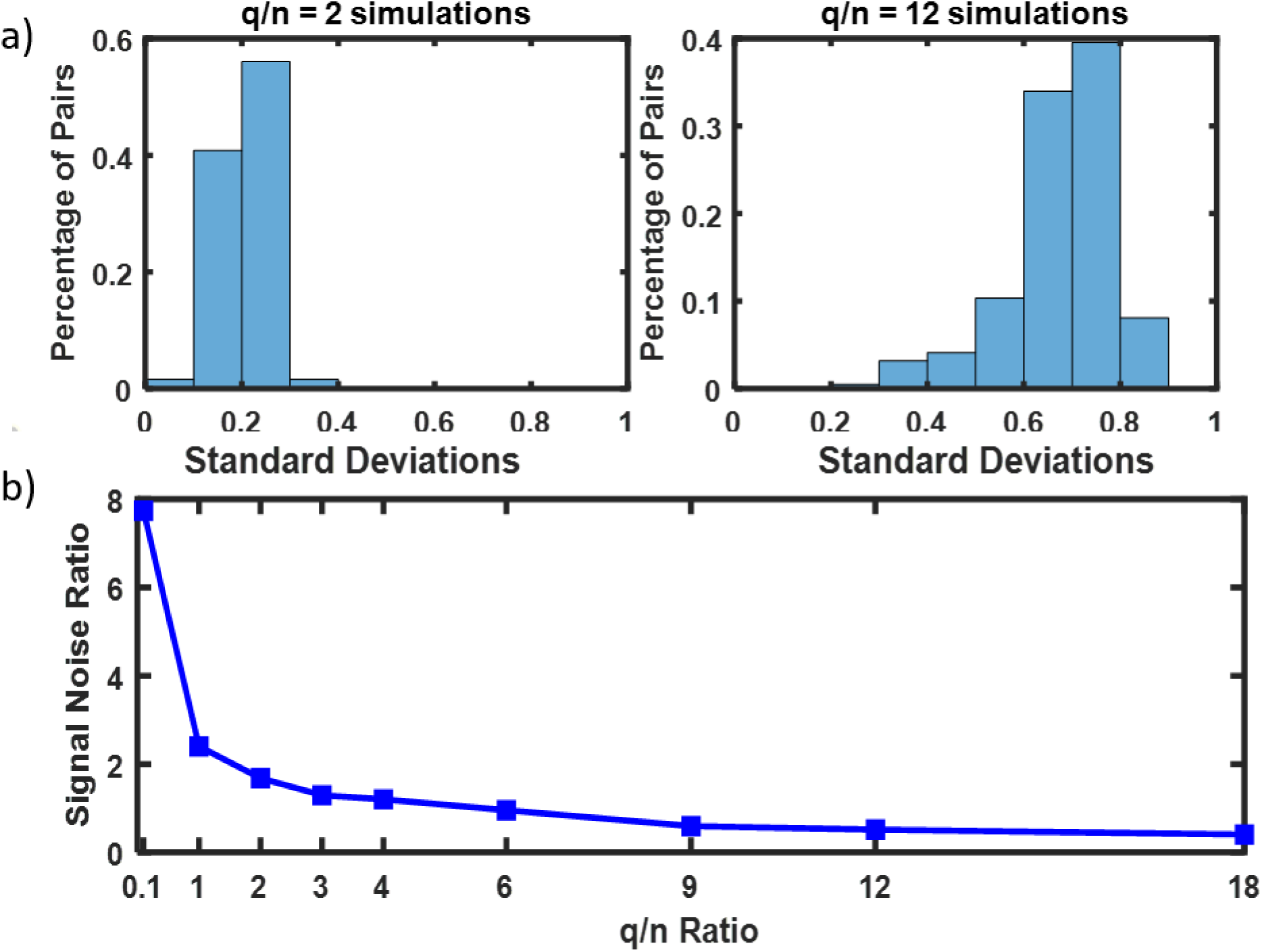
Explanation of increased variability of the simulated matrices. A- histograms of standard deviations of the elements of the simulated connectivity matrices for various q/n; B- signal to noise ratio vs. *q/n*.

#### Statistical Tests

The statistical tests for differences across clusters in this paper include Chi-square tests (MATLAB^®^ function crosstab) for categorical data, analysis of variance (ANOVA, MATLAB^®^ function anova1) for continuous data that follow a normal distribution, and the Kruskal-Wallis test (MATLAB® function kruskalwallis) for continuous data that do not follow a normal distribution. We controlled for the false discovery rate from multiple hypothesis testing using the Benjamini-Hochberg procedure (MATLAB^®^ function mafdr).

## Results and discussion

Here we demonstrate the results of cluster analysis of the four data sets described above by using the snakes-&-dragons approach. In clustering brain connectivity matrices from the 37 young and old healthy subjects pilot data set and the GSP data set, we provide not only the results of clustering but also the comparison with existing methods of correlation matrix comparison (RS, T-, and S-statistics), and evaluation of the quality of clustering. The microbiome example serves to illustrate the use of the dragon concept and demonstrates the Dragon 3 vector described above. The World Bank example demonstrates the broadness of the snakes-&-dragons approach and its applicability outside of the biomedical field.

### Brain connectivity matrices. Conventional measures vs. clustering of the snakes

The pilot data set of brain connectivity matrices of young and old healthy subjects was first used to examine the existing methods of matrix comparison. Pairwise distances between 37 brain connectivity matrices were determined by using RS, T-, and S-statistics. Then, hierarchical clustering was performed using the pairwise distances. The resulting dendrograms are presented in Fig 6; Fig 6a presents clustering based on RS, 6b on T-statistics, and 6c on S-statistics, while Fig 6d presents the results of hierarchical clustering of snake vectors. Dendrograms differ for the above four approaches, although all of them define two large clusters. Assuming that the true cluster membership is determined by the age of the participants, with 20 old participants and 17 young, we can calculate confusion matrices (Fig 6e-6h) as well as the misclassification error rate (Table 1) for each of the dendrograms. Note that the misclassification error is the lowest when the snake vector approach is used. Interestingly, however, the snake vector approach clustered three older brains (#10, 12, and 16) into the younger brain group, while all 17 young brains were correctly clustered together (Fig 6d). Notably, the use of random skewers also resulted in clustering of these three brains into the younger group (Fig 6a), while the use of the T-statistic clustered brain #10 into the younger group, and using the S-statistic clustered both brains #10 and #16 into the younger group. The problem with clustering real data is that one never knows the true class membership. Given the consensus between the four methods with regard to brain #10 and the consensus of three methods with regard to brain #16, it is possible that these brains preserved the properties of the young brains due to genetic or lifestyle factors despite their older age.

**Fig 6.**
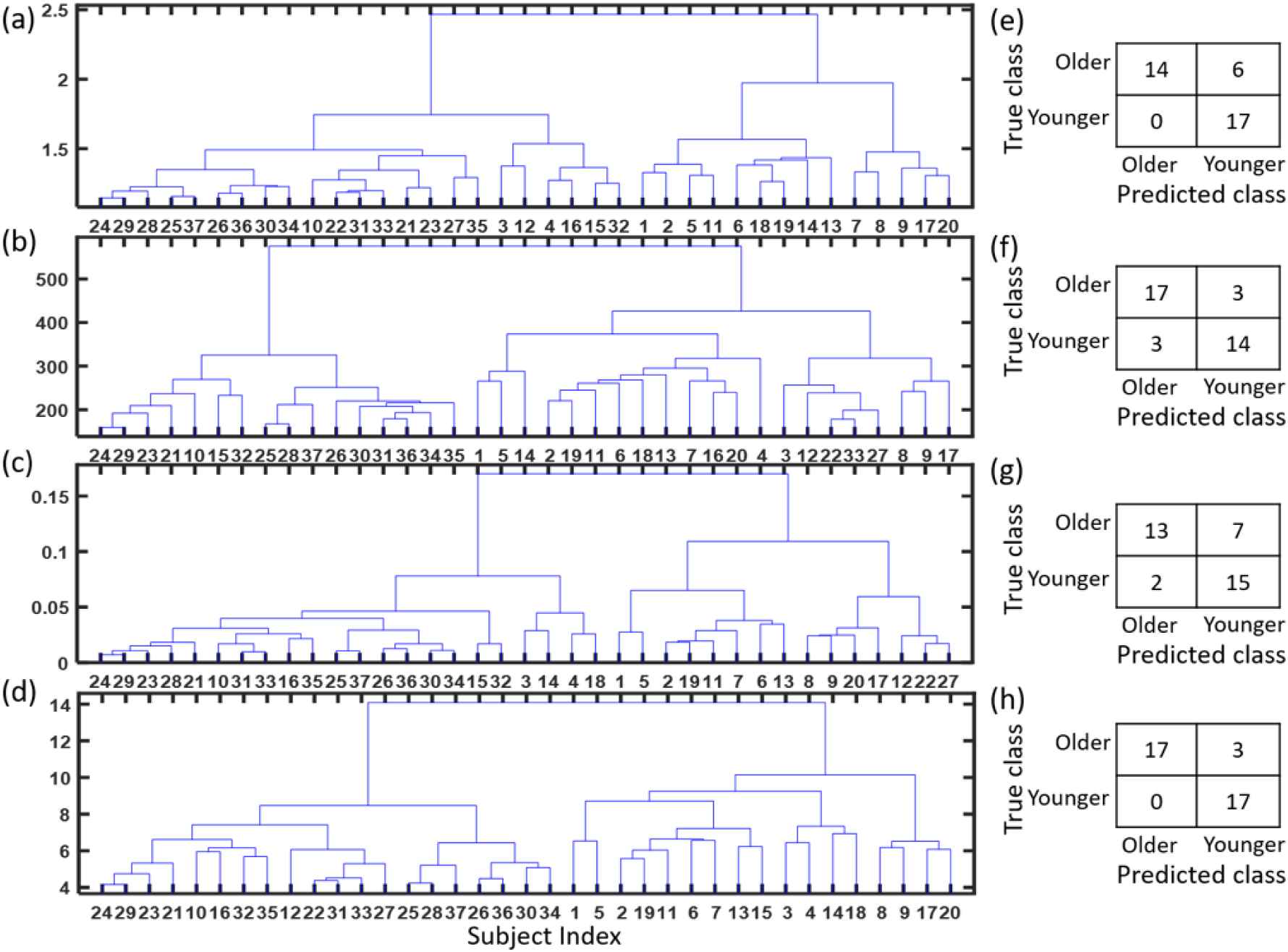
Clustering of brain connectivity matrices from pilot data set of young vs. old healthy persons. A-dendrogram based on RS, B-dendrogram based on T-statistics, C-dendrogram based on S-statistics, D-dendrogram based on snake vectors, E-H- confusion matrices for the above four approaches.

**Table 1.** Misclassification error of four clustering approaches in the pilot data set of brain connectivity matrices of young and old healthy subjects.

| Method | Old Group | Young Group | Misclassification Error |
| --- | --- | --- | --- |
| True Demographics | 20 | 17 | -- |
| Random Skewers + Hierarchical Clustering | 14 | 23 | 16.22% |
| T-statistic + Hierarchical Clustering | 21 | 16 | 13.51% |
| S-Statistic + Hierarchical Clustering | 15 | 22 | 24.32% |
| "Snake" Vector + Hierarchical | 17 | 20 | 8.10% |

In order to further evaluate the quality of clustering with the snakes approach, we used the simulated data created from the prototypical young (#29) and old (#1) brain connectivity matrices, as described in the Methods section. Note that brains #29 and #1 are distinctly different according to dendrograms from all four clustering methods (Fig 6). Since we know the true cluster memberships for the simulated data, we can calculate misclassification error for each clustering algorithm (Fig 7). Here in addition to using hierarchical clustering with RS, T- and S-statistics, and snake vectors, we examined the use of snake vectors with k-means clustering and with resampling-based consensus clustering (as described in Methods section). Misclassification errors up to *q/n* = 4 (SNR≥1.203 as defined by eq. 1) are all zero for all methods. For *q/n* > 6 (SNR<0.956), clustering correlation matrices using snake vectors outperforms the existing methods by having the lowest misclassification error rates, regardless of the clustering method used. The best performance is demonstrated by consensus clustering of snake vectors due to higher robustness to the added random noise.

**Fig 7.**
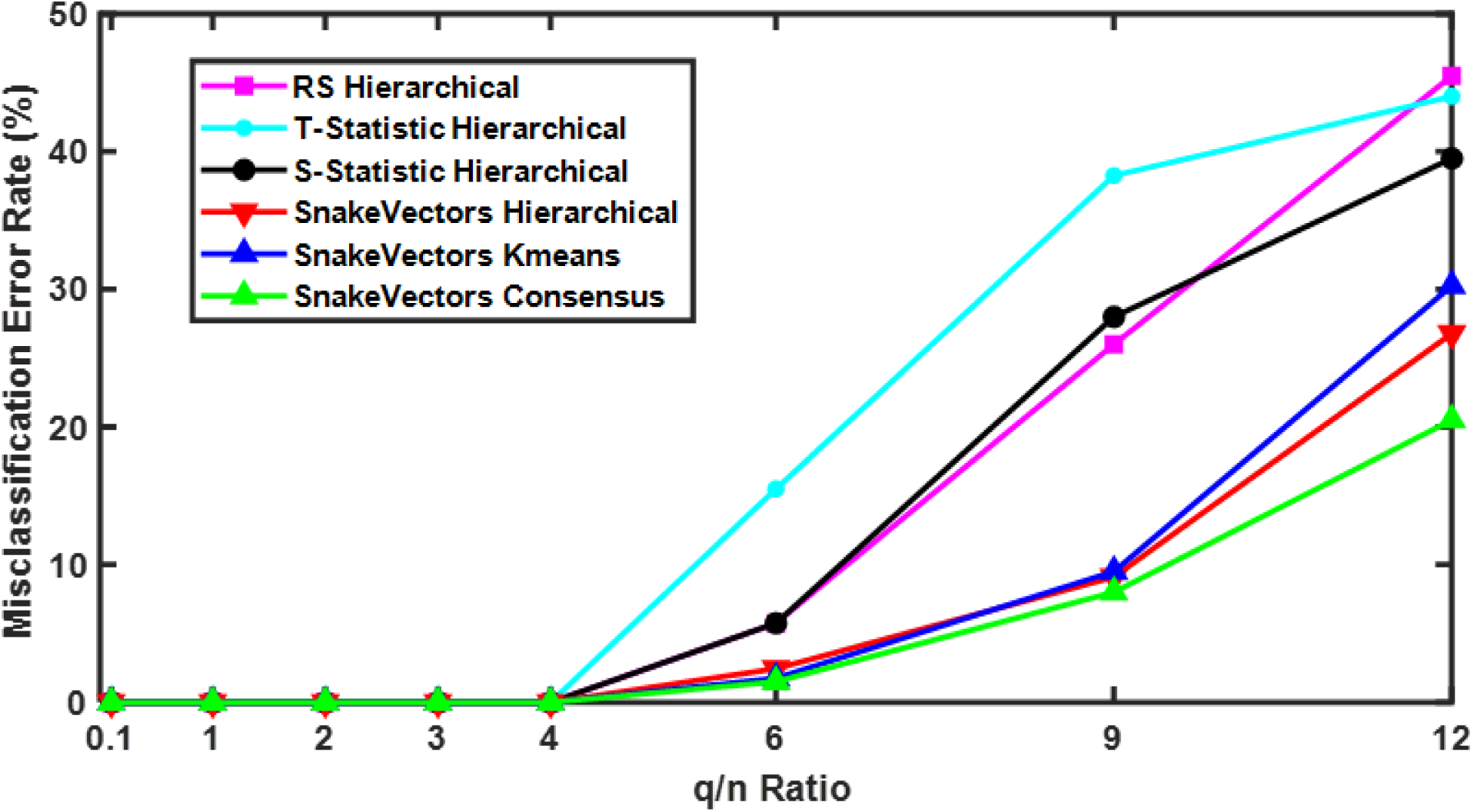
Misclassification error in clustering simulated connectivity matrices. Comparison of hierarchical clustering results for RS, T- and S-statistics, and snakes vectors, with k-means and resampling-based consensus clustering using snake vectors. Snake vectors based approaches outperform RS, T- and S-statistics based ones.

### Clustering of 500 brain connectivity matrices from the GSP project

Next, we applied our snake vectors approach to the clustering of 500 brain connectivity matrices from the GSP project. To cluster snake vectors derived from the connectivity matrices we used the resampling-based consensus clustering method as described in the Methods section. Fig 8a presents the heat map for the 500 x 500 consensus matrix. Each element of the matrix provides the probability that two brain connectivity matrices belong to the same cluster. Consensus clustering identified two distinct clusters with sample sizes N1= 160 and N2=340. Use of the Calinski criterion also confirmed the number of clusters as two (Fig 8b).

**Fig 8.**
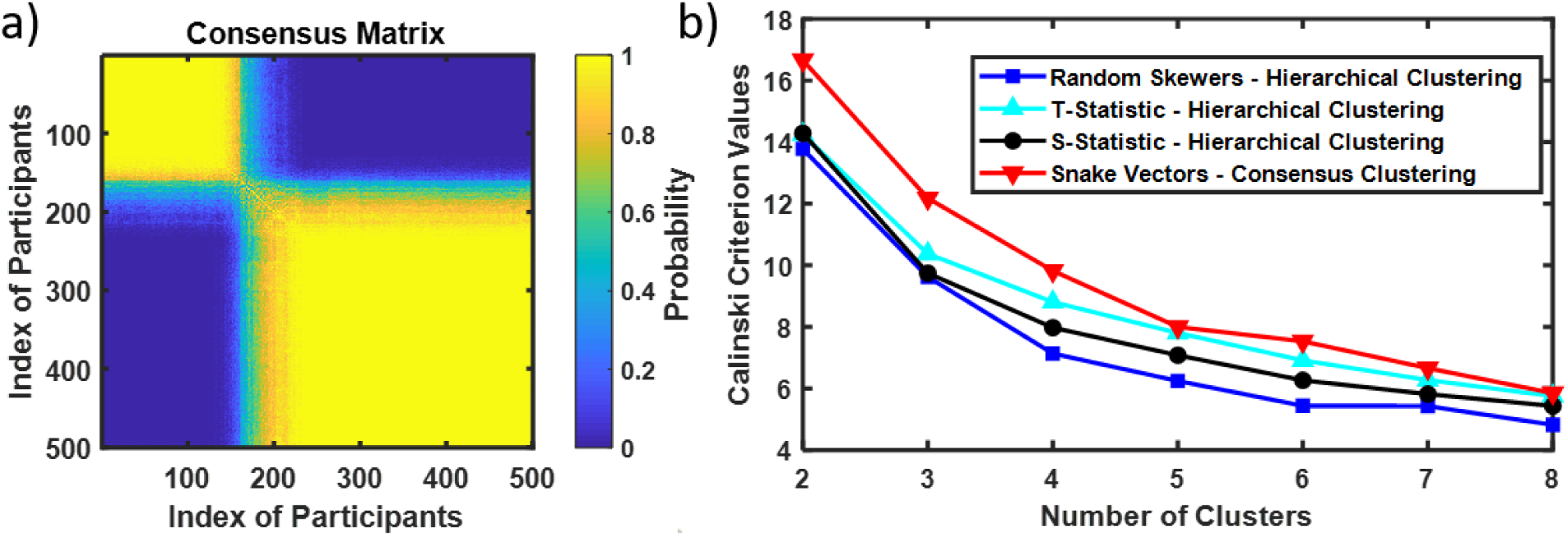
Resampling-based consensus clustering of 500 brain connectivity matrices from GSP project. A- Consensus matrix. Two identified clusters are presented as yellow squares (yellow color indicating the high probability of a pair of brains belonging to the same cluster). High contrast in the on diagonal and off-diagonal values of probability indicate two clusters. B- Checking the number of clusters with Calinski criterion. Calinski criterion have a maximum at k=2 indicating two clusters as well (both with snakes-&-dragons approach and with RS, T- and S-statistics).

Table 2 presents some anatomical and demographic variables of interest describing GSP participants but not used for clustering. Eight out of 81 such variables were significantly different across the two clusters; two of the variables remained significantly different after the correction for multi- testing (FDR corrected p-values < 0.05) [34]. Ethnicity was significantly different (FDR corrected p-value = 0.004) between the two clusters and sex was borderline significant (FDR corrected p-value = 0.055 and uncorrected p-value = 0.006), with cluster 2 having more white and female participants. Right vs. left handedness was not significant (*p=0.9*).

**Table 2.** Anatomical and demographic variables of interest describing GSP participants but not used for clustering

|  | Cluster 1<br>(n =160 ) | Cluster 2<br>(n = 340) |  |  |
| --- | --- | --- | --- | --- |
| Variables |  |  | p-value | FDR<br>corrected p |
| Age | 21.113(±2.63) | 21.335(±2.79) | 0.304 | 0.607 |
| Race/ethnicity |  |  | <0.001 | 0.004 |
| White not Hispanic | 83 (51.9%) | 233 (68.5%) |  |  |
| Other | 77 (48.1%) | 107 (31.5%) |  |  |
| Sex |  |  | 0.006 | 0.055 |
| Female | 81 (50.6%) | 216 (63.5%) |  |  |
| Male | 79 (49.4%) | 124 (36.5%) |  |  |
| Education | 14.231(±1.73) | 14.400(±1.72) | 0.234 | 0.575 |
| Handness |  |  | 0.906 | 0.947 |
| Right | 145 (91.2%) | 304 (89.9%) |  |  |
| Left | 14 (8.8%) | 34 (10.1%) |  |  |
| Right superior frontal thickness (mm) | 2.768(±0.13) | 2.798(±0.12) | 0.005 | 0.047 |
| Estimated total intracranial volume (cm<sup>3</sup>) | 1558.487(±146.8) | 1533.709(±140.0) | 0.027 | 0.191 |
| Right hemisphere average cortical thickness (mm) | 2.499(±0.07) | 2.514(±0.08) | 0.027 | 0.191 |
| Left hemisphere hippocampal volume (mm<sup>3</sup>) | 4490.225(±428.8) | 4420.709(±411.2) | 0.028 | 0.191 |
| Right hemisphere hippocampal volume (mm<sup>3</sup>) | 4511.075(±446.0) | 4441.971(±411.9) | 0.037 | 0.231 |
| Left inferiorparietal thickness (mm) | 2.434(±0.12) | 2.455(±0.11) | 0.04 | 0.232 |

Even more interesting is the comparison across the clusters of the variables that were used for clustering, i.e., the elements of the connectivity matrices. Fig 9a presents the average connectivity matrix for cluster 1 and Fig 9b for cluster 2. Fig 9c provides mean differences between connectivity matrices averaged across brains in cluster 2 and brains in cluster 1, while Fig 9d indicates by black dots which of the differences were significant (FDR corrected p-value < 0.05). A total of 8395 (out of 14196) elements of the connectivity matrices were significantly different even after the FDR correction for multi-testing [34]. Importantly, most of the significantly different elements of the connectivity matrices were not randomly distributed; they are rather concentrated within known brain subnetworks (defined in the Methods section and Fig 9 caption). Average correlation within the default mode network is significantly and substantially (over 26%) higher in cluster 2 than cluster 1, while the motor network is 26% more highly correlated in cluster 1 than cluster 2. Multiple average correlations between the known subnetworks were significantly different (FDR corrected p-value < 0.05) between cluster 1 and 2 as well, as shown in Table 3, e.g., VFN and CON are almost 215% more correlated in cluster 2 than in cluster 1. Importantly, the use of the snake vector approach allows identification of these distinctly different clusters.

**Fig 9.**
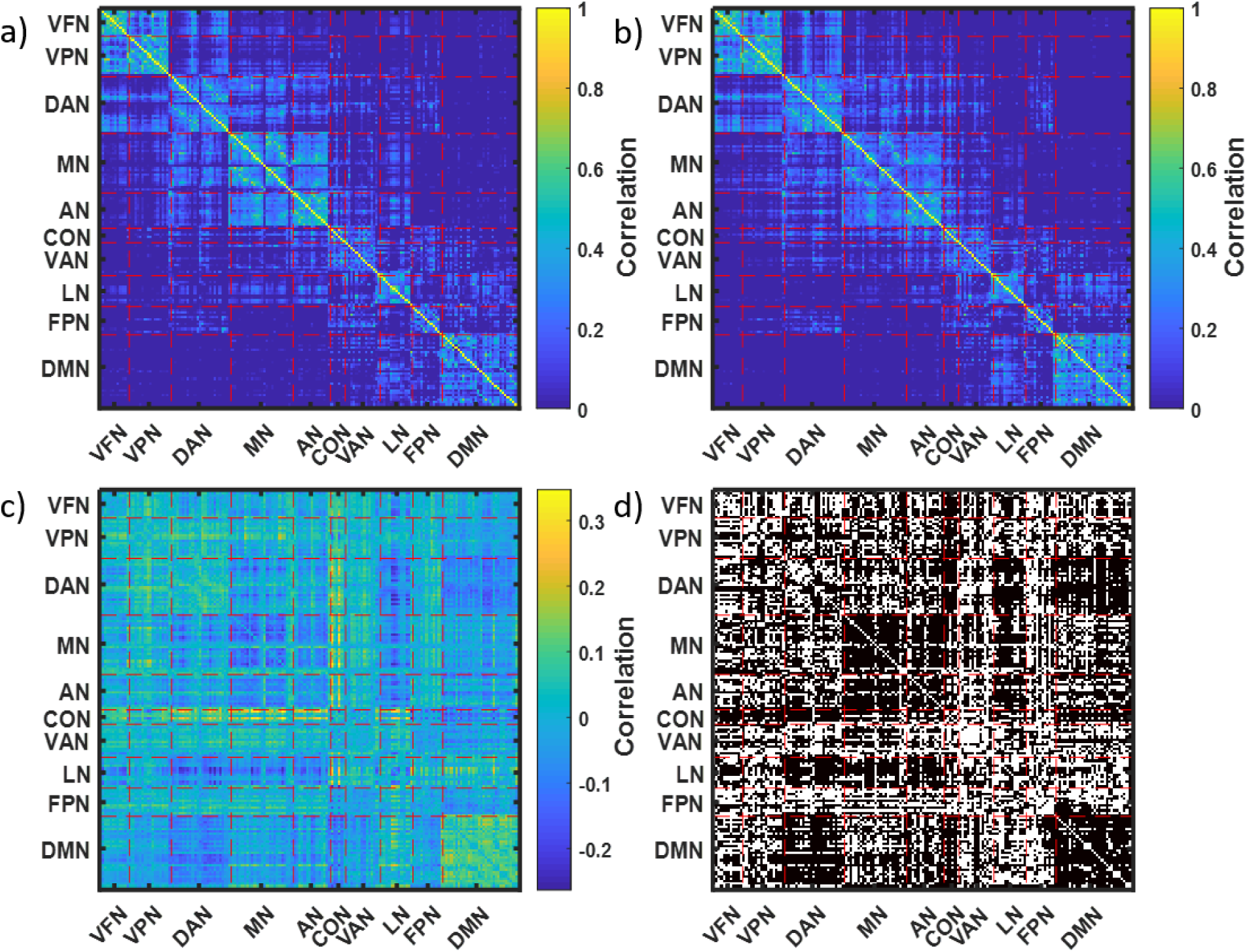
Mean brain connectivity matrices for two clusters identified in GSP data. A- Mean connectivity matrix for cluster 1, B- Mean connectivity matrix for cluster 2, C- Difference of mean connectivity matrices for cluster 2 and cluster 1, D- 8395 significantly different values of connectivity observed in cluster 1 vs. cluster 2. The 169 brain areas were divided into 10 networks: visual foveal (VFN), visual peripheral (VPN), dorsal attention (DAN), motor (MN), auditory (AN), cingulo-opercular (CON), ventral attention (VAN), language (LN), fronto-parietal (FPN), and default mode (DMN) [26].

**Table 3.**
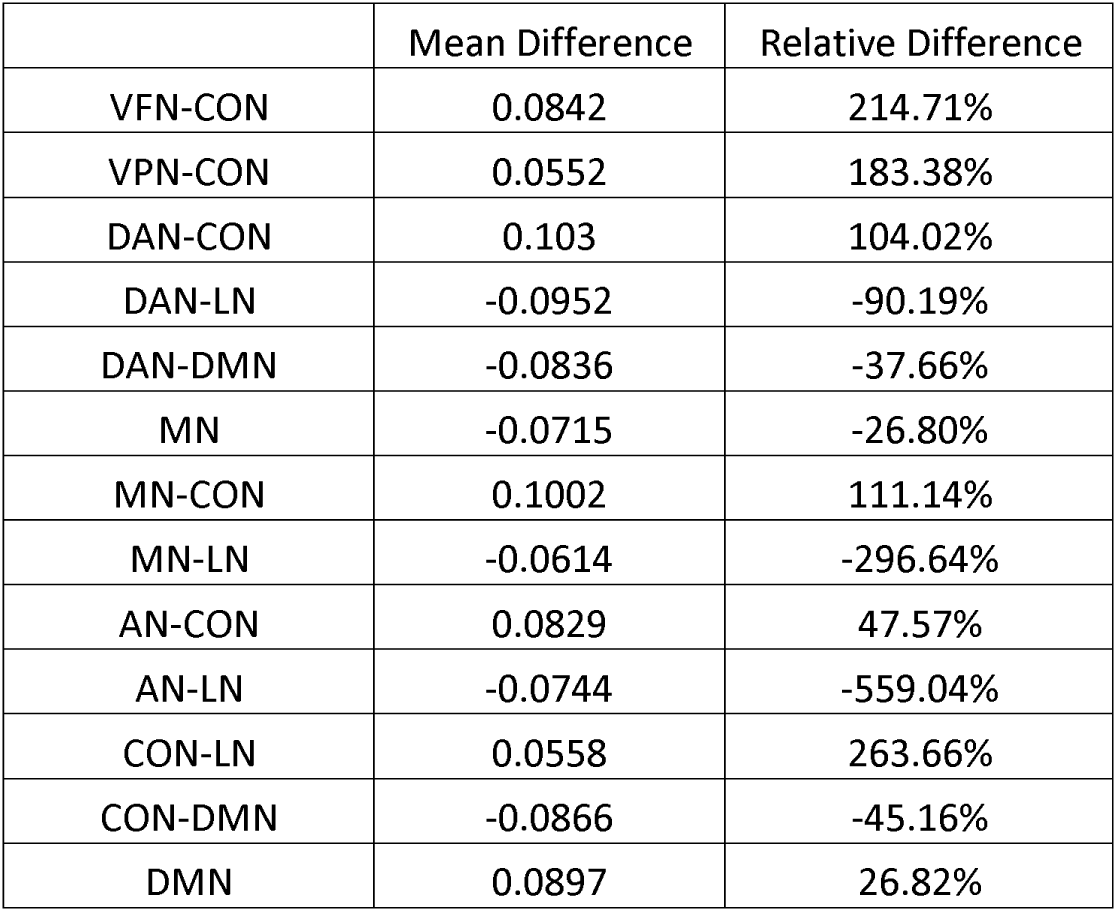
Significant differences in brain connectivity matrices are located mostly in the below subnetworks. Mean Difference: *c*_2_−*c*_1_. Relative Difference: *R*= (*c*_2_−*c*_1_)/*c*_1_, where c_1_ and c_2_ are the values of connectivity (correlation coefficients) averaged across the subnetworks in cluster 1 and cluster 2.

|  | Mean Difference | Relative Difference |
| --- | --- | --- |
| VFN-CON | 0.0842 | 214.71% |
| VPN-CON | 0.0552 | 183.38% |
| DAN-CON | 0.103 | 104.02% |
| DAN-LN | -0.0952 | -90.19% |
| DAN-DMN | -0.0836 | -37.66% |
| MN | -0.0715 | -26.80% |
| MN-CON | 0.1002 | 111.14% |
| MN-LN | -0.0614 | -296.64% |
| AN-CON | 0.0829 | 47.57% |
| AN-LN | -0.0744 | -559.04% |
| CON-LN | 0.0558 | 263.66% |
| CON-DMN | -0.0866 | -45.16% |
| DMN | 0.0897 | 26.82% |

### Using snakes-&-dragons for clustering of microbiomes of healthy college-age adults

For the microbiome data described in [13] and briefly in the Methods section, we calculated the correlations across OTU counts observed at seven time points (weeks) at four body sites (gut, tongue, palm, and forehead) to explore the temporal changes in each subject’s microbiome. We created 7×7 correlation matrices for each person and each body site to represent the similarities between the observed seven weeks in terms of the microbiome composition. We then conducted a cluster analysis using these correlation matrices and our snake vectors approach to identify subgroups of individuals sharing similar patterns of microbiome changes over time. We used three approaches to compare the above correlation matrices: 1) we clustered individuals by using data only from the gut and explored the correlation matrices for the other three sites; 2) we clustered the individuals using data from the gut, tongue, palm, and forehead separately; 3) we created dragon vectors by concatenating snake vectors for the gut, tongue, palm, and forehead and then clustered these dragon vectors. Analyses were performed on 52 students (out of 85 total) who provided samples from all four body sites for at least seven consecutive weeks. Figs 10-11 present the correlation matrices averaged across the members of the identified clusters. Note that students were clustered not by the composition of their microbiome, but rather by the pattern of change of their microbiomes over time, i.e., the dynamics of their microbiomes.

**Fig 10.**
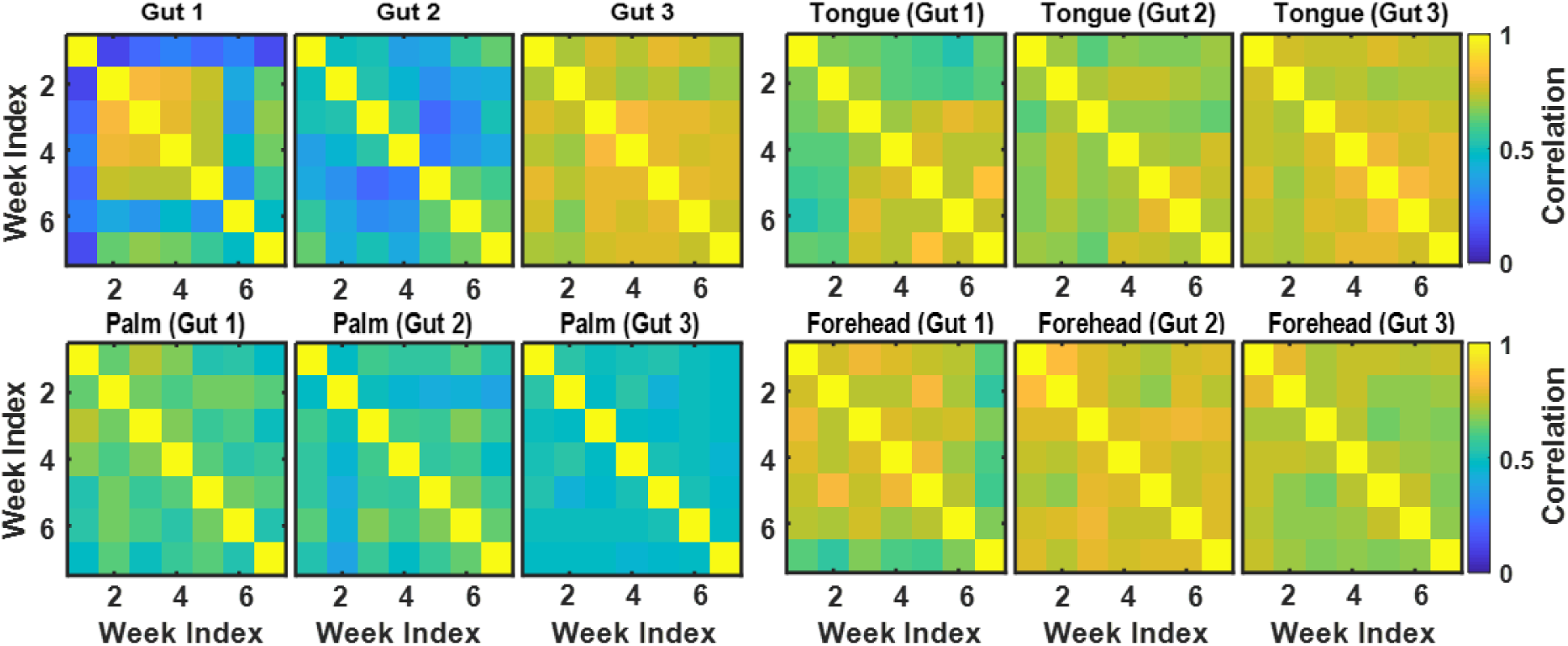
Correlation matrices reflecting microbiome dynamics at four body sites (gut, tongue, palm, and forehead) for three clusters of students identified based on the gut microbiome data.

**Fig 11.**
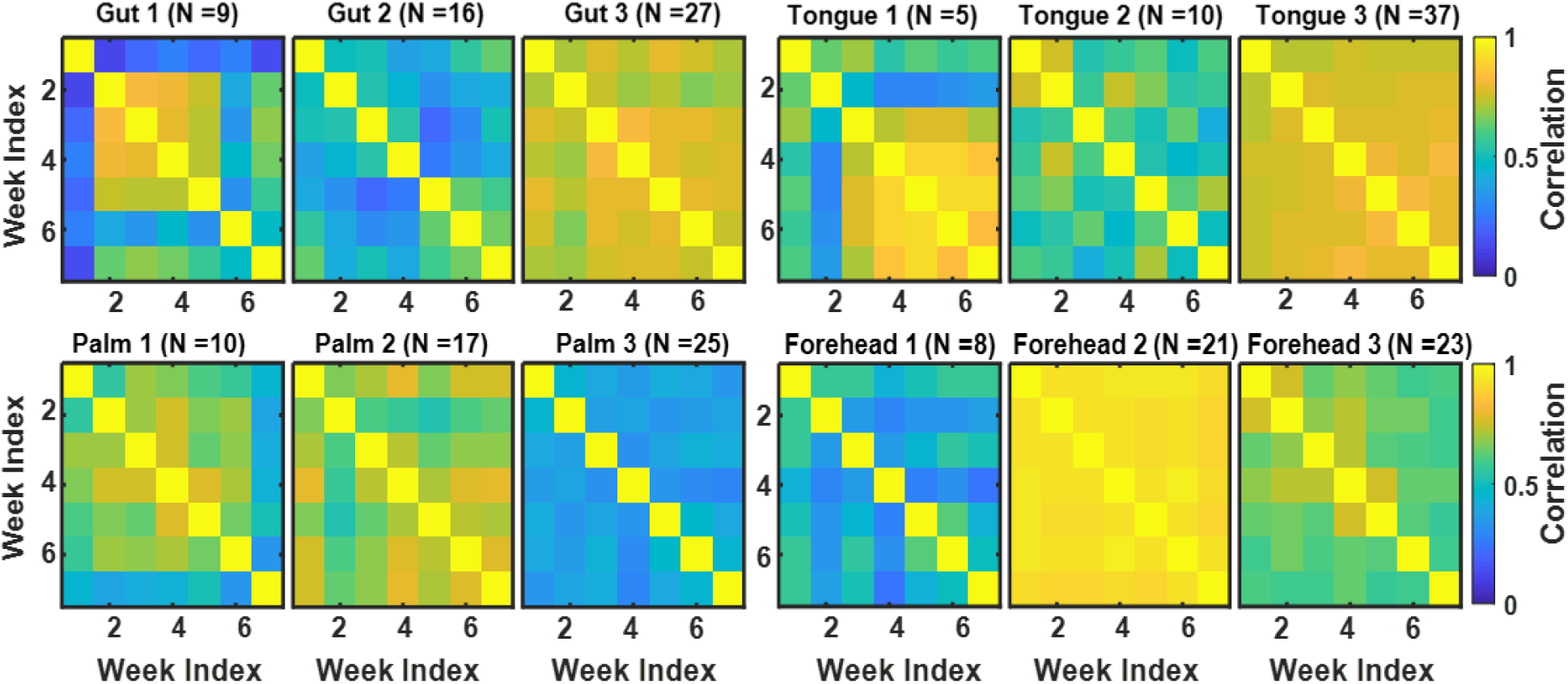
Correlation matrices reflecting microbiome dynamics at four body sites (gut, tongue, palm, and forehead) for three clusters of students identified based on the microbiome data for each of the body sites.

Fig 10 illustrates the first approach, where clustering is based on gut microbiome data, which resulted in three clusters named Gut 1 (n=9), Gut 2 (n=16), and Gut 3 (n=27). As seen in Fig 10a, for students in cluster Gut 1, the gut microbiome was highly correlated during weeks 2 through 5, while at weeks 1 and 6 their microbiomes were quite different from other weeks. There seems to have been some abrupt changes in the gut microbiomes of these students during weeks 1 and 6. For students in cluster Gut 2, the gut microbiome was moderately correlated across all 7 weeks and the level of correlation between the adjacent weeks was slightly oscillating in time. Students in cluster Gut 3 had stable gut microbiomes that did not change much over time. Comparison of the correlation matrices of tongue, palm, and forehead microbiomes for the Gut 1, 2, and 3 clusters (Figs 10b-10d) demonstrates that forehead and tongue microbiomes were relatively stable over time for all gut-based clusters, while the palm microbiome was less correlated over time. This is not surprising since palm microbiome communities are most affected by the environment in daily life.

In the second analysis, we clustered individuals based on the data from each of the four sites separately. The correlation matrices for each site averaged across each cluster are shown in Fig 11. We have identified three clusters in each of the four sites. Among these three clusters for each site, we have one cluster that has generally large correlation across all the weeks and one cluster that has relatively small correlation across all the weeks. We also have one or two clusters for each site that has one or two weeks that are quite different from the others; it is most pronounced in Gut 1, but is also present in Palm 1, Palm 2, Forehead 1, and Tongue 1. These peculiar weeks vary from site to site, which demonstrates different dynamics of the temporal evolution of microbial communities over the seven weeks.

Fig 12 provides Sankey diagrams for pairwise comparison of cluster membership across the four body sites. Note that cluster membership was similar when clustering was based on gut and tongue microbiomes—the most similar clusters being Gut 3 and Tongue 3.

**Fig 12.**
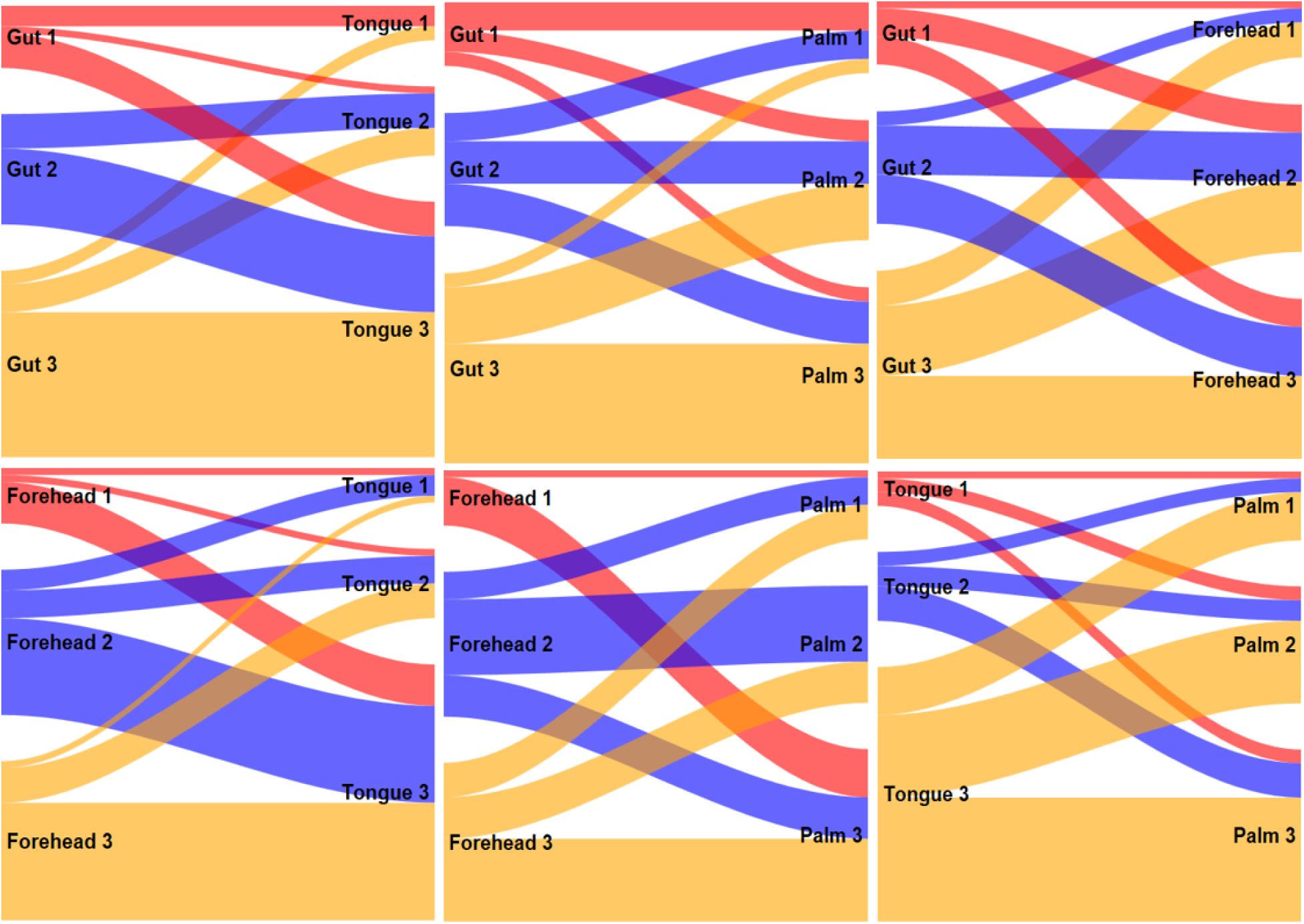
Pairwise comparison of cluster membership across four body sites.

In the third analysis, we clustered individuals using data from all four sites together. For each individual, we concatenated snakes from each site (forehead, tongue, gut, and palm) to form a “dragon” vector. We found three clusters: Body 1, 2, 3 (Fig 13a) with 12, 18, and 22 subjects in each cluster. For cluster Body 1, only the tongue microbiomes were highly correlated over time. For cluster Body 2, both tongue and gut microbiomes were highly correlated, while only the forehead microbiome was highly correlated over time for cluster Body 3. These results suggest the existence of subtypes representing different dynamics of microbial communities throughout the body. Sankey diagrams (Fig 13b) demonstrate that the cluster Body 3 is similar in membership to Forehead 2 and is driven by the high temporal stability of the forehead microbiome in this cluster. Cluster Body 2 is mostly formed by the members of the Tongue 3 cluster with highly stable tongue microbiome, and cluster Body 1 includes members of various site-specific clusters.

**Fig 13.**
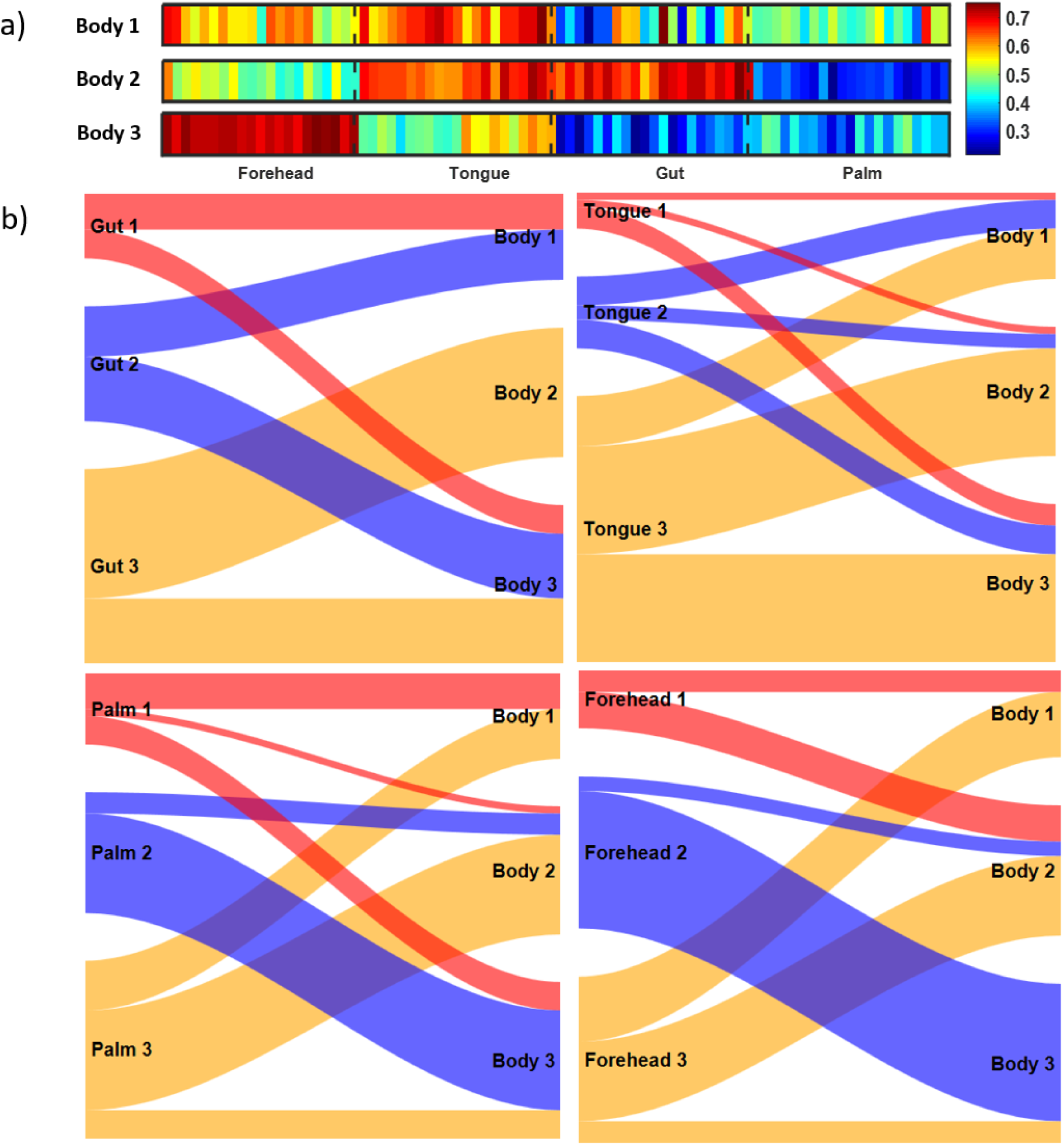
Clustering based on dragon vectors describing microbiomes of four body sites. A-Mean dragon vectors for three clusters of students identified by clustering the concatenated snake vectors for gut, tongue, palm, and forehead. B-Sankey diagrams comparing cluster membership based on the dynamics of microbiomes at each site and all four sites’ microbiomes combined.

Table 4 provides overall microbiome, demographic, and behavioral data for each of the clusters identified in the above analyses, allowing interpretation and providing possible reasons for the similarities and differences in the patterns of microbiome dynamics. Note that the actual microbiomes within the clusters could be quite different while the patterns of microbiome dynamics are similar. The top three rows of the table characterize the diversity of the microbiome within the given site averaged across the members of each cluster. The total number of OTUs (which can serve as one of the measures of microbiome diversity) was calculated by counting the OTUs that were observed in a sample from any week for each student and then averaged across all the students in the given cluster and rounded to the closest integer. Each OTU was counted only once even if it was observed at multiple weeks. Another important measure of diversity is the Shannon Index (SI), defined as 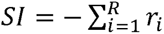 ln *r_i_*, where *r_i_* is the measure of relative abundance of the given OTU, i.e., the ratio of the abundance of the given OTU to the abundance of all observed OTUs, and *R* is the total number of observed OTUs for the given sample. The values of the SI for each student, site, and week from the supplementary data of [13] were averaged across the weeks and across the members of the identified clusters. The SI characterizes the diversity of the microbiome by taking into account not only the number of OTUs but their abundances as well [35]. Higher values of the index describe diverse populations; lower values of the index describe populations dominated by a single taxon (OTU). In the case of a single taxon, SI=0, while in the case of all taxa (OTUs) being represented equally SI= ln(R). In order to simplify the comparison of sites and students with different numbers of OTUs, we also calculated the normalized SI equal to SI/ln(R), which has the maximum possible value of one and minimum of zero.

**Table 4.** Overall microbiome, demographic, and behavioral data for each of the identified clusters based on dynamics of a) gut microbiome, b)tongue microbiome, c)palm microbiome, d)forehead microbiome, e) four sites microbiome.

| Gut-based clusters |  |  |  |  |  |
| --- | --- | --- | --- | --- | --- |
| Variables | Gut 1 (n=9) | Gut 2 (n=16) | Gut 3 (n=27) | p value | Corrected p |
| **Number of OTUs** | 969 | 1053 | 1042 | 0.2632 | 0.3948 |
| **Shannon Index** | 4.687 | 5.125 | 5.275 | 0.014 | 0.0652 |
| **Normalized Shannon Index** | 0.817 | 0.871 | 0.883 | 0.029 | 0.0652 |
| **Age** | 20.778 | 25.438 | 23.962 | 0.1416 | 0.2549 |
| **BMI** | 22.915 | 22.446 | 23.077 | 0.7402 | 0.7737 |
| **Gender** |  |  |  | 0.7468 | 0.7737 |
| Female | 6 (67%) | 10 (63%) | 14 (54%) |  |  |
| Male | 3 (33%) | 6 (37%) | 12 (46%) |  |  |
| **Race /Ethnicity** |  |  |  | 0.0148 | 0.0652 |
| Caucasian | 4 (44%) | 14 (93%) | 22 (81%) |  |  |
| Hispanic | 1 (11%) | 1 (7%) | 3 (11%) |  |  |
| Other | 4 (44%) | 0 (0%) | 2 (7%) |  |  |
| **University** |  |  |  | 0.0251 | 0.0652 |
| UCB | 6 (67%) | 6 (38%) | 14 (52%) |  |  |
| NAU | 0 (0%) | 5 (31%) | 12 (44%) |  |  |
| NCS | 3 (33%) | 5 (31%) | 1 (4%) |  |  |
| **Use of Facial Cosmetics** |  |  |  | 0.7737 | 0.7737 |
| Never | 4 (44%) | 5 (31%) | 9 (33%) |  |  |
| Rarely | 0 (0%) | 3 (19%) | 4 (15%) |  |  |
| Occasionally | 1 (11%) | 1 (6%) | 0 (0%) |  |  |
| Regularly | 1 (11%) | 1 (6%) | 2 (7%) |  |  |
| Daily | 3 (33%) | 6 (38%) | 12 (44%) |  |  |
| Tongue-based clusters |  |  |  |  |  |
| Variables | Tongue 1 (n=5) | Tongue 2 (n=10) | Tongue 3 (n=37) | p value | Corrected p |
| **Number of OTUs** | 364 | 380 | 326 | 0.1945 | 0.3501 |
| **Shannon Index** | 3.424 | 4.002 | 4.156 | 0.0015 | 0.0135 |
| **Normalized Shannon Index** | 0.819 | 0.800 | 0.699 | 0.0033 | 0.0146 |
| **Age** | 21.000 | 23.000 | 24.459 | 0.3152 | 0.4301 |
| **BMI** | 25.878 | 23.613 | 22.231 | 0.1121 | 0.2522 |
| **Gender** |  |  |  | 0.9759 | 0.9759 |
| Female | 3 (60%) | 5 (56%) | 22 (59%) |  |  |
| Male | 2 (40%) | 4 (44%) | 15 (41%) |  |  |
| **Race /Ethnicity** |  |  |  | 0.3345 | 0.4301 |
| Caucasian | 3 (60%) | 8 (80%) | 29 (81%) |  |  |
| Hispanic | 0 (0%) | 1 (10%) | 4 (11%) |  |  |
| Other | 2 (40%) | 1 (10%) | 3 (8%) |  |  |
| **University** |  |  |  | 0.0440 | 0.1320 |
| UCB | 5 (100%) | 4 (40%) | 17 (46%) |  |  |
| NAU | 0 (0%) | 2 (20%) | 15 (41%) |  |  |
| NCS | 0 (0%) | 4 (40%) | 5 (14%) |  |  |
| **Use of Facial Cosmetics** |  |  |  | 0.8882 | 0.9759 |
| Never | 1 (20%) | 4 (40%) | 13 (35%) |  |  |
| Rarely | 1 (20%) | 2 (20%) | 4 (11%) |  |  |
| Occasionally | 0 (0%) | 0 (0%) | 2 (5%) |  |  |
| Regularly | 1 (20%) | 0 (0%) | 3 (8%) |  |  |
| Daily | 2 (40%) | 4 (40%) | 15 (41%) |  |  |

| Palm-based clusters |  |  |  |  |  |
| --- | --- | --- | --- | --- | --- |
| Variables | Palm 1 (n=10) | Palm 2 (n=17) | Palm 3 (n=25) | p value | Corrected p |
| **Number of OTUs** | 1552 | 1648 | 2063 | 0.0656 | 0.1969 |
| **Shannon Index** | 5.449 | 5.533 | 6.099 | 0.0288 | 0.1801 |
| **Normalized Shannon Index** | 0.896 | 0.898 | 0.968 | 0.04 | 0.1801 |
| **Age** | 24.200 | 22.688 | 24.480 | 0.2662 | 0.4278 |
| **BMI** | 22.294 | 22.414 | 23.364 | 0.8090 | 0.8090 |
| **Gender** |  |  |  | 0.1114 | 0.2507 |
| Female | 7 (70%) | 6 (37%) | 17 (68%) |  |  |
| Male | 3 (30%) | 10 (63%) | 8 (32%) |  |  |
| **Race / Ethnicity** |  |  |  | 0.5395 | 0.6069 |
| Caucasian | 6 (60%) | 15 (88%) | 19 (79%) |  |  |
| Hispanic | 2 (20%) | 1 (6%) | 2 (8%) |  |  |
| Other | 2 (20%) | 1 (6%) | 3 (13%) |  |  |
| **University** |  |  |  | 0.3488 | 0.4485 |
| UCB | 7 (70%) | 8 (47%) | 11 (44%) |  |  |
| NAU | 1 (10%) | 5 (29%) | 11 (44%) |  |  |
| NCS | 2 (20%) | 4 (24%) | 3 (12%) |  |  |
| **Use of Facial Cosmetics** |  |  |  | 0.2852 | 0.4278 |
| Never | 3 (30%) | 6 (35%) | 9 (36%) |  |  |
| Rarely | 2 (20%) | 0 (0%) | 5 (20%) |  |  |
| Occasionally | 1 (10%) | 1 (6%) | 0 (0%) |  |  |
| Regularly | 1 (10%) | 0 (0%) | 3 (12%) |  |  |
| Daily | 3 (30%) | 10 (59%) | 8 (32%) |  |  |
| Forehead-based clusters |  |  |  |  |  |
| Variables | Forehead 1 (n=8) | Forehead 2 (n=21) | Forehead 3 (n=23) | p value | Corrected p |
| **Number of OTUs** | 1772 | 1771 | 1465 | 0.0579 | 0.1042 |
| **Shannon Index** | 5.595 | 4.077 | 5.609 | <0.0001 | 0.0001 |
| **Normalized Shannon Index** | 0.9022 | 0.8993 | 0.6704 | <0.0001 | <0.0001 |
| **Age** | 22.875 | 24.600 | 23.565 | 0.6866 | 0.7724 |
| **BMI** | 21.174 | 22.954 | 23.305 | 0.3887 | 0.4998 |
| **Gender** |  |  |  | 0.0048 | 0.0144 |
| Female | 8 (100%) | 7 (35%) | 15 (65%) |  |  |
| Male | 0 | 13 (65%) | 8 (35%) |  |  |
| **Race / Ethnicity** |  |  |  | 0.1488 | 0.2233 |
| Caucasian | 4 (50%) | 16 (80%) | 20 (87%) |  |  |
| Hispanic | 1 (12%) | 2 (10%) | 2 (9%) |  |  |
| Other | 3 (38%) | 2 (10%) | 1 (4%) |  |  |
| **University** |  |  |  | 0.9101 | 0.9101 |
| UCB | 5 (63%) | 9 (43%) | 12 (52%) |  |  |
| NAU | 2 (25%) | 8 (38%) | 7 (30%) |  |  |
| NCS | 1 (13%) | 4 (19%) | 4 (17%) |  |  |
| **Use of Facial Cosmetics** |  |  |  | 0.0361 | 0.0811 |
| Never | 4 (50%) | 5 (24%) | 9 (39%) |  |  |
| Rarely | 3 (38%) | 1 (5%) | 3 (13%) |  |  |
| Occasionally | 0 (0%) | 1 (5%) | 1 (4%) |  |  |
| Regularly | 1 (13%) | 0 (0%) | 3 (13%) |  |  |
| Daily | 0 (0%) | 14 (67%) | 7 (30%) |  |  |

| Body (Four body sites-based clusters) |  |  |  |  |  |
| --- | --- | --- | --- | --- | --- |
| Variables | Body 1 (n=12) | Body 2 (n=18) | Body 3 (n=22) | p value | Corrected p |
| **Number of OTUs** | 3551 | 3627 | 3221 | **0.0324** | 0.0728 |
| **Shannon Index** | 4.899 | 5.01 | 4.68 | **0.0017** | **0.0153** |
| **Normalized Shannon Index** | 0.602 | 0.612 | 0.581 | **0.0066** | **0.0207** |
| **Age** | 21.917 | 24.944 | 24.048 | 0.4636 | 0.5961 |
| **BMI** | 23.912 | 22.589 | 22.585 | 0.6881 | 0.7311 |
| **Gender** |  |  |  | **0.0069** | **0.0207** |
| Female | 10 (83%) | 13 (72%) | 7 (33%) |  |  |
| Male | 2 (17%) | 5 (28%) | 14 (66%) |  |  |
| **Race /Ethnicity** |  |  |  | 0.7311 | 0.7311 |
| Caucasian | 9 (75%) | 13 (72%) | 18 (86%) |  |  |
| Hispanic | 1 (8%) | 3 (17%) | 1 (5%) |  |  |
| Other | 2 (17%) | 2 (11%) | 2 (9%) |  |  |
| **University** |  |  |  | 0.1639 | 0.2459 |
| UCB | 7 (58%) | 9 (50%) | 10 (45%) |  |  |
| NAU | 1 (8%) | 8 (44%) | 8 (36%) |  |  |
| NCS | 4 (33%) | 1 (6%) | 4 (18%) |  |  |
| **Use of Facial Cosmetics Use** |  |  |  | 0.1145 | 0.2061 |
| Never | 5 (42%) | 7 (39%) | 6 (27%) |  |  |
| Rarely | 2 (17%) | 4 (22%) | 1 (5%) |  |  |
| Occasionally | 1 (8%) | 0 (0%) | 1 (5%) |  |  |
| Regularly | 2 (17%) | 2 (11%) | 0 (0%) |  |  |
| Daily | 2 (17%) | 5 (28%) | 14 (64%) |  |  |

As noted in [13], the highest diversity in terms of the number of OTUs and the highest SI values were observed at the skin surfaces (palm and forehead) which are most exposed to contacts with the environment. However, the highest values of SI and normalized SI of all skin sites were observed for Palm 3 (SI=6.10) and Forehead 3 (SI=5.61), which demonstrated low correlation of microbiomes across the 7 weeks. The microbiomes of the forehead-based clusters were significantly affected by the use facial cosmetics (p-value 0.036), e.g., Forehead 2 is characterized by the highest percentage (67%) of members using facial cosmetics daily, relatively low value of SI=4.08, and high value of normalized SI =0.9, indicating nearly equal representation of all OTUs.

Gut-based and tongue-based clusters demonstrated lower diversity in terms of lower numbers of OTUs, and lower SI and normalized SI values. The lowest values of the Shannon Index were observed in Tongue 1 (SI=3.42) and Gut 1 (SI=4.69), which also demonstrated abrupt changes in microbiomes at least twice in 7 weeks. The important role of the Shannon Index in predicting stability of the microbiome was already discussed in [13]; here we confirm this observation for the sites less exposed to environmental influences and identify clusters of participants with lower gut and tongue microbiome stability, which also demonstrated lower microbiome diversity. The explanation for lower diversity or stability of the microbiome in these groups of students is not clear. It might be related to race and ethnicity since the less stable clusters Gut 1 and Tongue 1 have a higher proportion of non-Caucasians and non-Hispanics (reported as race/ethnicity=other in Table 4). These clusters also have a higher proportion of students from the University of Colorado, Boulder and may be hypothetically related to some of them eating at the same places (e.g., school cafeterias). It is possible that the lower diversity and stability is caused by the actual composition of the microbiomes and its evolution over time, analysis of which would require construction of the covariance matrices (and snakes-&-dragons) not across weeks, but across OTUs, which will be the focus of our next paper. Nevertheless, having the ability to group individuals by microbiome variability instead of microbiome composition may prove to be a powerful tool in identifying disease predilection especially given the personalized nature of the human microbiome [36–37]. Future studies could also leverage our tool using case-control studies of disease with known microbiome components to determine if temporal groupings have health relevance.

### Clustering snakes based on macroeconomics development indicators from the World Bank

To demonstrate the use of the snake vectors approach outside of the biomedical field, we created 7×7 correlation matrices for economies of 200 countries using annual data collected by the World Bank. In particular, we looked at seven important macroeconomic indices: 1) gross domestic product (GDP); 2) unemployment; 3) inflation; 4) net trade in goods; 5) labor force participation; 6) foreign direct investment; and 7) gross domestic savings. Fig 14 illustrates the results of clustering of these correlation matrices using our snake vectors approach. Each of the presented matrices are the average of the correlation matrices of the above seven macroeconomic indices across the economies belonging to the given cluster. We also fit linear regression models to assess the amount of variability (R^2^) in 170 other development indicators that could be explained by the eight cluster groups. Among those with highest R^2^ was annual GDP growth, which had a significant (p<0.001) association with the eight cluster groups and therefore may help to elucidate the different mechanisms that can drive economic growth. For example, cluster 6 had high positive correlations between GDP and unemployment, yet had the highest growth. Although initially unexpected, this result may inform novel strategies and new macroeconomic models for economic growth in developing countries such as India, Mongolia, and Egypt, all of which were in cluster 6. Thus, clustering on correlations between macroeconomic indicators may identify novel subgroups representing different economic structures.

## Conclusions

We presented a novel method named “snakes-&-dragons” for comparing and subtyping of complex systems through clustering of vectors derived from the correlation matrices of the variables describing these systems. Using a real dataset and a simulated dataset on brain connectivity matrices, we showed that the novel approach outperformed the existing methods for comparison of correlation matrices (RS, T-, and S-statistics). In the analysis of brain connectivity matrices from the GSP project, our approach allowed identification of two clusters with distinctly different patterns of brain connectivity not explained by differences in demographic variables. In the analysis of the microbiome of healthy students, it allowed identification of clusters of students with distinctly different patterns of microbiome dynamics. It also allowed formulation of the hypothesis that stability of gut and tongue microbiomes is affected by the diversity of the microbiome (as described by the Shannon Index). The macroeconomic example illustrated the possibility of using the snakes-&-dragons approach outside of the biomedical field.

**Fig 14.**
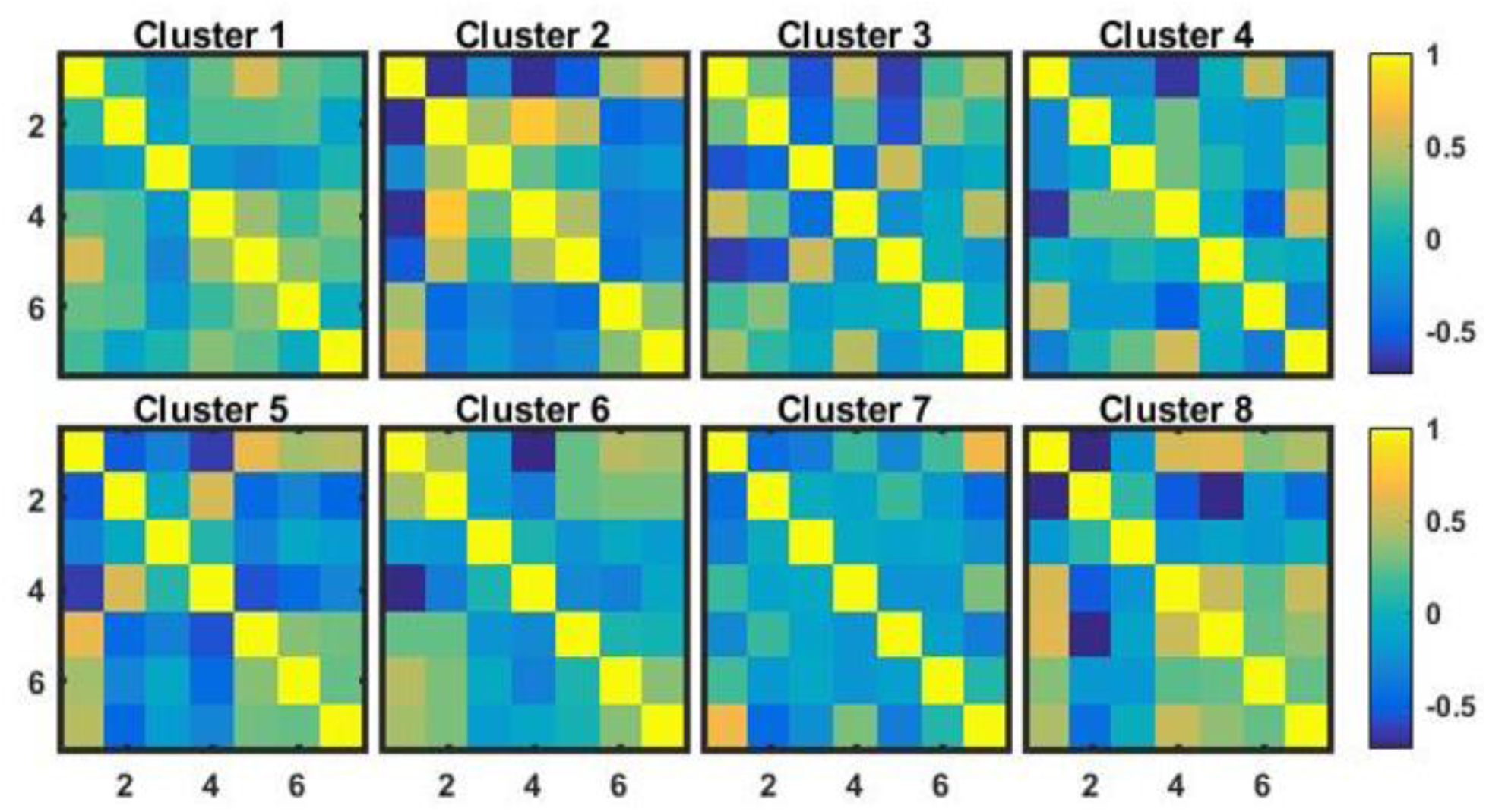
Correlation matrices of macroeconomic indices of eight identified clusters of economies.

We have developed a clustering method capable of unsupervised classification of objects based on their structures and interactions of their parts and attributes, therefore uncovering new patterns/groupings based on previously unexplored characteristics of the systems. In medicine, it could lead to identification of new, more homogeneous subtypes of complex common diseases and subsequently to more targeted treatments. As for limitations, we have not yet demonstrated all of the capabilities of the dragon vectors. For instance, in the analysis of the microbiome data it would be meaningful to combine in a dragon vector the snake vectors formed from the correlation matrices across the weeks and the correlation matrices across the OTUs. In drug discovery, it would be informative to combine correlation matrices formed from the multidimensional time series of transcriptomics and proteomics data collected at various time points after the perturbation of a cell culture with the drugs of interest. We plan to explore these capabilities in our future research.

A reader of this paper may be inclined to ask, “Does it really matter how to form a snake vector, or is it just about forming a vector that includes all the elements of the upper triangle of the correlation matrix?” Our answer to this question evolved from “Not really” to “Yes and No”, and eventually to “Well, yes”, and is worth explaining here. If there is no intrinsic order of the variables upon which correlations are calculated, then the order in which correlation matrices and snakes are formed does not matter; it is important, however, that the order of variables should be the same in all correlation matrices under comparison and that the order of correlation coefficients used in snake formation should be the same as well. Similarly, the concatenation of multiple snakes or other data elements in the formation of dragons should be consistent across objects. In case an intrinsic order of variables does exist, the situation is different. Take, for instance, the situation where different time points are compared as in our microbiome example; in this case, the first “off-diagonal” of the matrix demonstrates the correlations between measurements separated by one week, the second “off- diagonal” separated by two weeks, etc. Creating snakes in any other way than the serpentine of “off- diagonals” would violate this natural order. Imagine now the situation where the system has “memory” of limited duration (such as in a Markov process); in this case, the correlation matrix would look like a ribbon of nonzero elements along the diagonal and several “off-diagonals” with zeros everywhere else, so the snake vectors representing such matrices could be truncated. Another case of intrinsic order is physical distance. We believe that the snake vector approach could be useful in analysis of Hi-C data [38–40], where the conformation of DNA in the chromosomes is derived from the matrix of distances between the nucleotides or larger elements of genome. In this case, the intrinsic variable is the distance from the beginning of the DNA chain. The periodicity of the elements of the snake vectors constructed as an off-diagonal serpentine would be informative of the DNA conformation. These matrices are huge, so the truncation of the snake vectors that represent them are computationally beneficial when possible. Even more interesting is the situation where the intrinsic order is distance in 3D space, e.g., the distance from the tumor or a lesion to the multiple locations in which biomarkers are measured. In this case, a higher dimensional analog of a correlation matrix is required which should be described by objects more complex than snakes-&-dragons, bringing to mind creatures like Zmey Gorynych from Russian folk tales – a dragon with 3 heads [41].

## Acknowledgements

Data were provided in part by the Brain Genomics Superstruct Project of Harvard University and the Massachusetts General Hospital (Principal Investigators: Randy Buckner, Joshua Roffman, and Jordan Smoller), with support from the Center for Brain Science Neuroinformatics Research Group, the Athinoula A. Martinos Center for Biomedical Imaging, and GSP Open Access Documentation the Center for Human Genetic Research. Twenty individual investigators at Harvard and MGH generously contributed data to the overall project.

The authors want to thank Dr. Shimony from Washington University for providing pre-processed connectivity matrices and for helpful discussions. We also thank the Washington University Alzheimer’s Disease Research Center for providing normative fMRI data.

